# Central vestibular tuning arises from patterned convergence of otolith afferents

**DOI:** 10.1101/2020.02.14.948356

**Authors:** Zhikai Liu, Yukiko Kimura, Shin-ichi Higashijima, David G. Hildebrand, Joshua L. Morgan, Timothy E. Holy, Martha W. Bagnall

## Abstract

As sensory information moves through the brain, higher-order areas exhibit more complex tuning than lower areas. Though models predict this complexity is due to convergent inputs from neurons with diverse response properties, in most vertebrate systems convergence has only been inferred rather than tested directly. Here we measure sensory computations in zebrafish vestibular neurons across multiple axes *in vivo*. We establish that whole-cell physiological recordings reveal tuning of individual vestibular afferent inputs and their postsynaptic targets. An independent approach, serial section electron microscopy, supports the inferred connectivity. We find that afferents with similar or differing preferred directions converge on central vestibular neurons, conferring more simple or complex tuning, respectively. Our data also resolve a long-standing contradiction between anatomical and physiological analyses by revealing that sensory responses are produced by sparse but powerful inputs from vestibular afferents. Together these results provide a direct, quantifiable demonstration of feedforward input convergence *in vivo.*

## Introduction

Neurons compute information from many different synaptic inputs. A central challenge in understanding neuronal circuits is determining how the tuning and connectivity of these inputs affect the resulting computations. For example, neurons in visual cortex exhibit simple or complex orientation tuning, which is thought to derive from the convergence of presynaptic inputs with distinct tuning properties (Hubel and Wiesel, 1962, Alonso and Martinez, 1998). Computational models of such input-output relationships have fundamentally shaped the way we think of information processing in the brain (Felleman and Van Essen, 1991, LeCun et al., 2015). However, these models generally require assumptions about many parameters that can only be measured with incompatible approaches: the tuning of the presynaptic population, input connectivity, and synaptic strengths, as well as the activity of the postsynaptic neuron itself. Direct measurements of these parameters simultaneously are prohibitively difficult in most systems, making it hard to define neuronal computations *in vivo*.

Vestibulospinal (VS) brainstem neurons receive direct vestibular sensory inputs from peripheral vestibular afferents (Boyle et al., 1992) and project to the spinal cord (Boyle and Johanson, 2003). Understanding the neuronal computations of VS neurons would not only inform how vestibular sensory signals are processed in the brain, but also provide a mechanistic view of sensorimotor transformation. VS neurons, like other central vestibular neurons, produce diverse responses to head movement. During head tilt or acceleration, some central vestibular neurons exhibit simple cosine-tuned responses, similar to those of the afferents: the strongest activity is evoked by movements in a preferred direction, with little or no response in the orthogonal direction. In contrast, other central vestibular neurons exhibit more complex responses, including bidirectional responses (Peterson, 1970) and spatiotemporally complex tuning (Angelaki et al., 1993). A vectorial model predicts that convergence of several simple cosine-tuned afferents can fully account for the response of either a simple or a complex central vestibular neuron, depending on whether those afferents are similarly tuned or differently tuned (Angelaki, 1992). However, as in other systems, this model has been technically challenging to test experimentally.

We chose to address this question in the small brain of the larval zebrafish. The VS circuit was previously identified in the larval zebrafish as the homolog of mammals (Kimmel et al., 1982), which becomes functional as early as 3 days post fertilization (dpf) (Mo et al., 2010). The accessibility of the larval zebrafish brain for intracellular recording from identified VS neurons allows us to investigate how central vestibular neurons compute sensory signals in vertebrates.

Here we establish a novel approach to record sensory evoked responses *in vivo* from VS neurons in the larval zebrafish. We find that individual afferents evoke large amplitude-invariant excitatory postsynaptic currents (EPSCs), allowing us to separate distinct afferent inputs that converge onto a given VS neuron. This provides a mechanism to simultaneously measure the sensory tuning and synaptic strength of each converging afferent, as well as the response of the postsynaptic neuron. We show that afferents with similar tuning direction preferentially converge, producing simple tuning in the VS neuron. Furthermore, the smaller number of cells with complex bidirectional responses receive input from differently tuned afferents, with consequent simple or complex spiking. We also show that these afferent inputs are sufficient to predict the tuning of the VS neuron. Together, this work reveals how central neurons in the brain compute sensory information from their presynaptic inputs.

## Results

### Sensory evoked responses of vestibulospinal neurons *in vivo*

Traditionally, measurements of neuronal responses to vestibular stimuli have been accomplished by unit recordings (Angelaki and Dickman, 2000, Schor et al., 1984, Fernandez and Goldberg, 1976a). Directly measuring vestibular-evoked synaptic currents in central neurons *in vivo* has been technically challenging (Arenz et al., 2008, Chabrol et al., 2015). We designed a custom whole-cell electrophysiology rig to deliver translational motion stimuli to 4-7 dpf larval zebrafish via an air-bearing motorized sled (Fig. 1A). This setup allows intracellular measurement of sensory-evoked responses from vestibulospinal (VS) neurons on multiple axes *in vivo* for the first time, to the best of our knowledge. To target identified VS neurons, we generated a Tg(nefma:gal4; UAS:GFP) line, whose labelling overlaps dye backfilling (Figs.1D, S1) from the spinal cord, consistent with evidence of *Nefm* expression in mammalian vestibular neurons (Kodama et al., 2020). We recorded spontaneous EPSCs in voltage clamp, at overall rates varying from 1 to 365 EPSC/s. Delivery of translational movement evoked corresponding modulations in EPSC frequency (Fig. 1B). The extent of modulation varied depending on the direction of the stimulus delivered across four different axes (Fig. 1C). In this example neuron, EPSC rate was modulated most strongly in the rostral-caudal (R-C) axis and weakly in the dorsal-ventral (D-V) axis, with intermediate strength responses for the diagonal directions (R/D-C/V and R/V-C/D).

**Figure 1:**
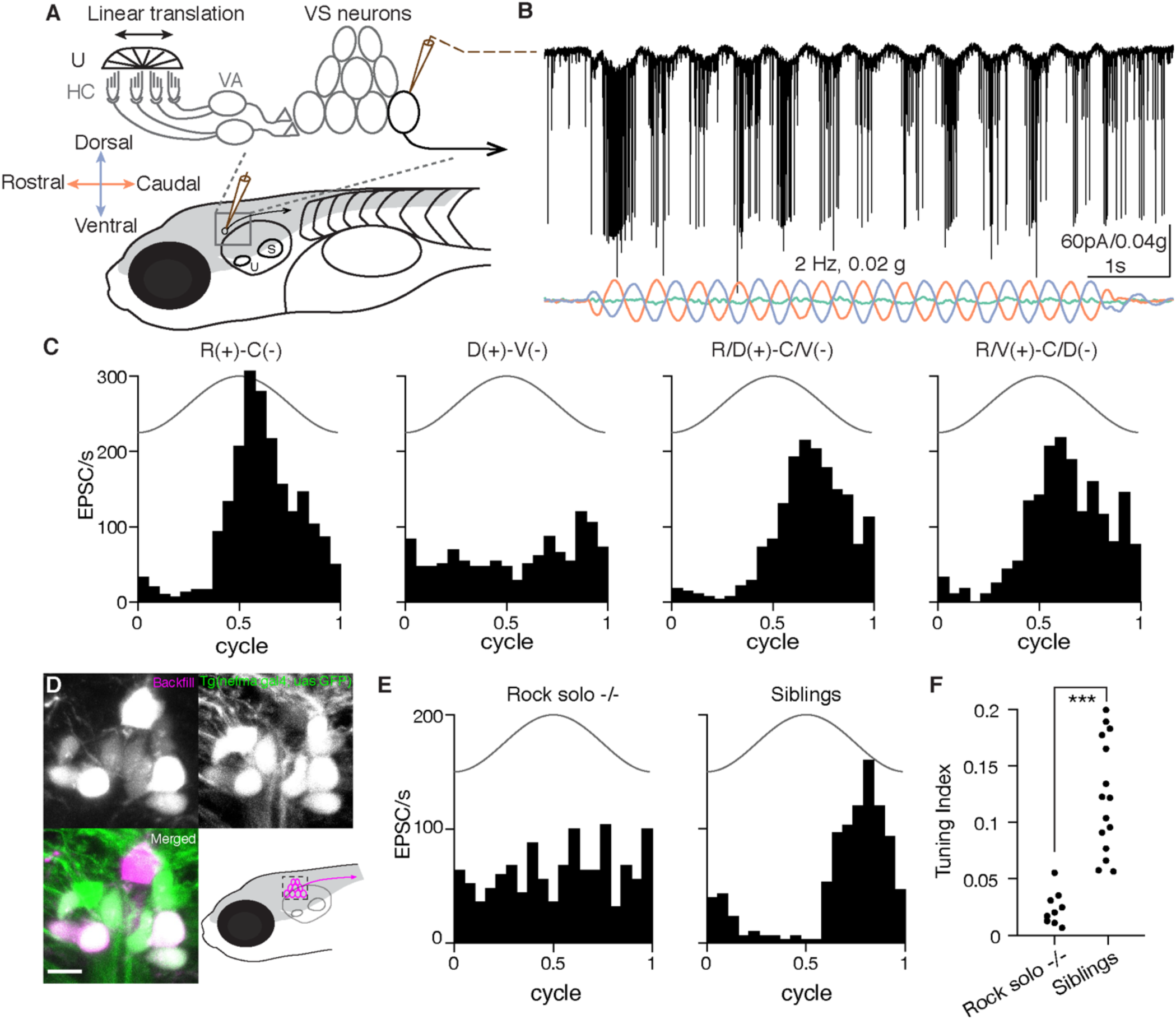
Sensory-evoked responses *in vivo* in vestibulospinal (VS) neurons. A. Schematic representing *in vivo* patch clamp recording configuration and vestibular afferent circuit in larval zebrafish, U: utricle, HC: hair cells, VA: vestibular afferents, S: saccule. B. Example recording trace from a VS neuron in voltage clamp mode during 2 Hz, 0.02 g translational movement on the R/V(+)-C/D(-) axis. EPSCs, black; acceleration in three body axes, colored (orange, rostral[+]-caudal[-]; green, dorsal[+]-ventral[-]; blue, ipsilateral[+]-contralateral[-]). C. Sensory-evoked EPSC responses to translation in four different axes for the same VS neuron as in B, across 12 cycles. Solid line, acceleration (2 Hz, 0.02 g). D. Tg(nefma:gal4; UAS:GFP) (green) colabels VS neurons identified by dye backfilling (magenta) from spinal cord. Scale bar: 5 *µ*m E. Sensory responses of a VS neuron in the best direction in a rock solo -/- (left) and in a het/WT sibling (right). F. Summary of tuning index in the best direction for all VS neurons recorded in rock solo -/- (9 neurons, 5 fish) and siblings (15 neurons, 10 fish). Mann-Whitney U test, p=6.5e-5

Response to translational movement could derive from the vestibular or other sensory inputs. In larval zebrafish, the anterior otolith (utricle) is the sole functional vestibular sensor (Riley and Moorman, 2000). To examine whether utricular signaling is necessary for the observed tuning, we measured the sensory response of VS neurons in the *otog*^*c.1522+2T>A*^ *-/-* (rock solo) animals, which lack the utricle (Mo et al., 2010, Roberts et al., 2017). Translational stimuli were ineffective at modulating EPSC rate in VS neurons of rock solo homozygotes in contrast to wild-type/heterozygous siblings (representative examples, Fig. 1E). Across all recordings, VS neurons of rock solo -/- animals exhibit largely untuned EPSCs compared to siblings, as quantified by a tuning index (Fig. 1F). Thus, this approach reveals sensory-evoked synaptic responses encoding directional vestibular stimuli in identified VS neurons *in vivo*.

### Mixed electrical and chemical synapses mediate the transmission from otolith afferents to VS neurons

What properties define the vestibular afferent synapse onto VS neurons? In rodents, vestibular afferent synapses onto vestibulo-ocular reflex neurons exhibit amplitude-invariant synaptic transmission, mediated by specialized vesicular release machinery (Bagnall et al., 2008, McElvain et al., 2015, Turecek et al., 2017). To characterize afferent synaptic input to VS neurons, we electrically stimulated the vestibular (anterior statoacoustic) ganglion while recording from VS neurons in voltage clamp (Fig. 2A). Stimulation evoked a synaptic current with two components. The first component had fast kinetics with short latency (0.56 ± 0.28 ms, n=8), low jitter (0.05 ± 0.04 ms, n=8), and invariant EPSC amplitude (SD: 6.7±3.9%, normalized to peak) across trials. In contrast, the second component had slower kinetics and variable amplitudes (Fig. 2B). We dissected the two components of evoked EPSCs pharmacologically. Bath application of the gap junction blocker carbenoxolone (CBX, 500 *µ*M) during afferent stimulation substantially reduced the first component of the EPSC (Figs. 2C, E). In contrast, bath application of the AMPA receptor antagonist NBQX (10 *µ*M) abolished the second component of synaptic current (Fig. 2D). Furthermore, the fast EPSCs were not reversed by changing the holding potential (Fig. S2), a signature behavior of electrical synaptic transmission (Akrouh and Kerschensteiner, 2013). Thus, the early and late components of afferent-evoked synaptic currents are mediated by gap junctions and AMPA receptors, respectively. Across VS neurons, the NBQX-sensitive currents accounted for 27.1±20.2% of total charge transfer (n=7, Fig. 2F), demonstrating that gap junctional current is the major component mediating synaptic transmission.

**Figure 2:**
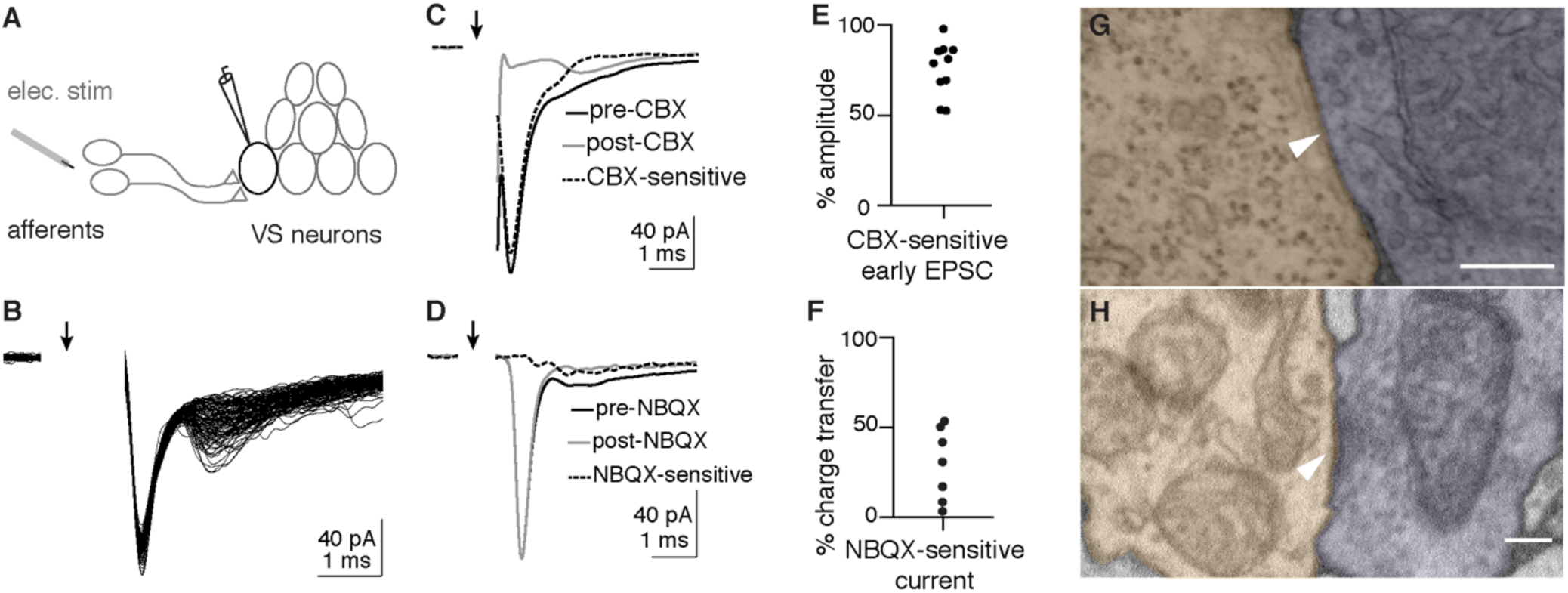
Otolith afferent to VS neuron transmission is mediated by mixed electrical and chemical synapses. A. Schematic of whole-cell recording configuration from VS neuron while electrically stimulating otolith afferents. B. Example EPSCs evoked by electrical stimulation of the otolith afferents; 105 EPSCs overlaid. Arrow indicates onset of stimulation. Stimulus artifact is blanked. C. Carbenoxolone (CBX) diminishes the fast component of evoked EPSCs D. NBQX abolishes the second, slower component of evoked EPSCs E. Group data quantifying the reduction of early EPSC amplitude by CBX F. Group data quantifying the total charge transfer that is abolished by NBQX application G. Example EM image of gap junction between identified otolith afferent (pseudocolored purple) and VS neuron (orange). Scale bar: 200 nm. H. Example EM image of chemical synapse between otolith afferent (purple) and VS neuron (orange). Scale bar: 200 nm.

To evaluate ultrastructural evidence for mixed synaptic transmission, we re-imaged existing serial ultrathin sections of a 5.5 dpf larval zebrafish (Hildebrand et al., 2017) at sufficiently high resolution (1-4 nm/px) to identify synaptic contacts between myelinated utricular afferents and VS neurons, identified anatomically. We found both tight junction structures (Fig. 2G), and vesicles apposed to a postsynaptic density (Fig. 2H) at appositions between utricular afferent and VS neurons, consistent with anatomical evidence for mixed electrical / chemical transmission at this synapse in adult fish (Korn et al., 1977) and rat (Nagy et al., 2013). Together, these results demonstrate that VS neurons receive vestibular afferent inputs mediated by amplitude-invariant gap junctional (electrical) and variable amplitude glutamatergic (chemical) synapses.

### Inferring afferent tuning from distinct EPSCs

Because electrically mediated EPSCs from afferents to VS neurons exhibited a fixed amplitude, we hypothesized that we could distinguish the activity of individual otolith afferents converging onto a given VS neuron by their characteristic EPSC amplitudes. Indeed, spontaneous and sensory-evoked EPSCs recorded in VS neurons often fell into distinctive size bins, as visualized in a histogram of EPSC amplitudes (Figs. 3A, B). EPSCs were sorted into three clusters with unsupervised learning (see Methods), primarily leveraging their amplitudes. Each of these EPSC clusters showed a stereotypical amplitude and waveform in this example neuron (Fig. 3A, inset). To test whether each cluster of EPSCs amplitudes corresponds to an individual afferent, we used an approach derived from spike sorting: temporal autocorrelation to test for refractory periods within EPSC event times. Physiologically, one afferent cannot generate two action potentials within its refractory period (Fernandez et al., 1972); thus the EPSCs elicited by that afferent should exhibit a refractory period as well. An auto-correlogram of all EPSC event times in this example neuron did not display a refractory period (Fig. 3C, top). In contrast, an auto-correlogram within each EPSC cluster exhibited a clear refractory period around 0 ms (Fig. 3C, bottom). Furthermore, cross-correlograms between EPSC clusters did not show this structure, consistent with the notion that they arise from independent inputs (Fig. S3). Accordingly, we can interpret these three EPSC clusters as deriving from the activity of three distinct presynaptic afferents (Fig. 3D, left). Because of the high fidelity of electrical transmission, each EPSC cluster effectively reads out the spiking of an individual afferent, allowing us to measure presynaptic activity via postsynaptic recording (Fig. 3D, right).

**Figure 3:**
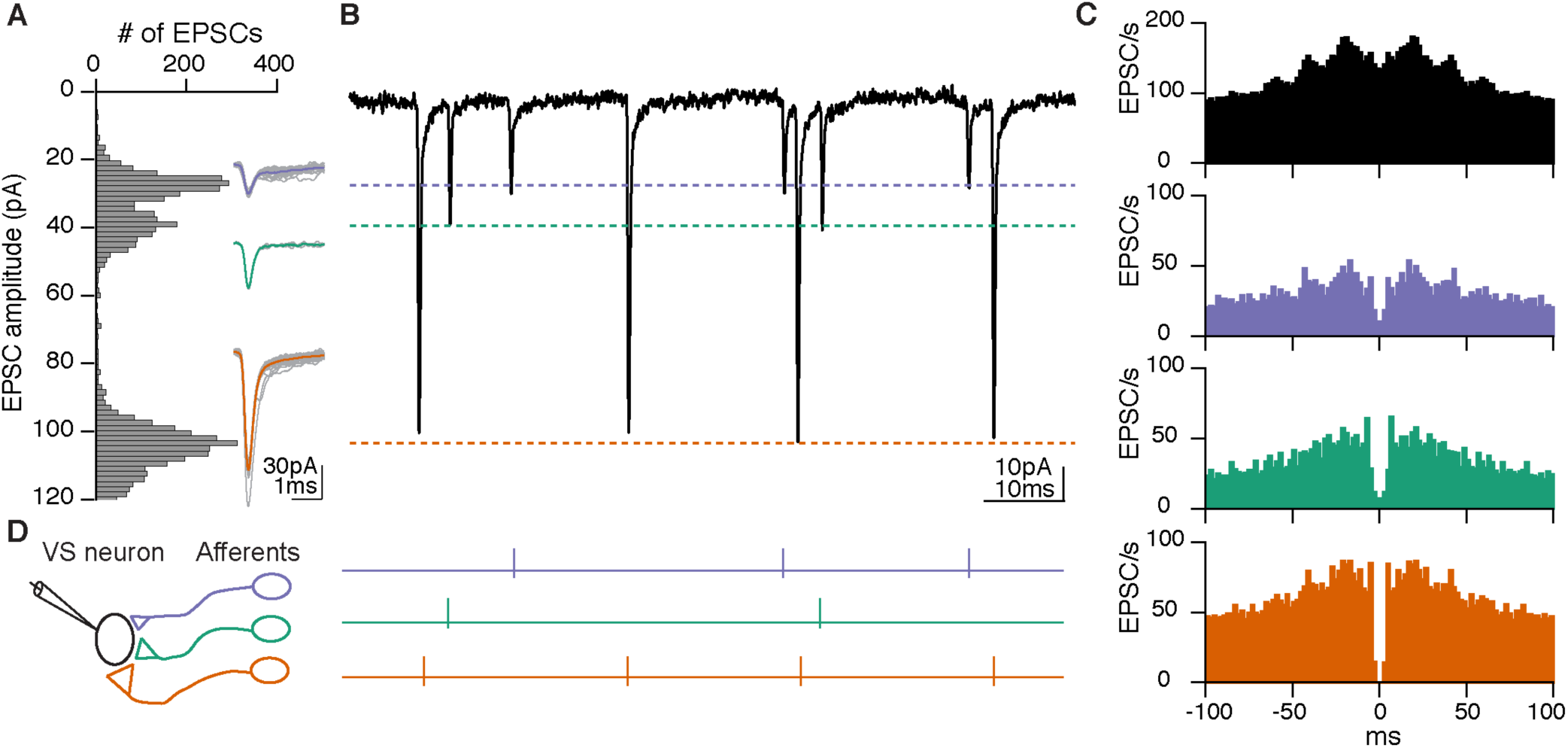
Distinct EPSC amplitudes reflect individual afferent inputs. A. Histogram of spontaneous and sensory-evoked EPSC amplitude distribution of the same VS neuron as Fig. 1B. Inset, overlay of individual EPSCs (gray) and average (colored) for each amplitude bin. B. Example trace of EPSCs exhibiting stereotypic shapes and amplitudes in three clusters, corresponding to each amplitude bin in A. C. Auto-correlogram of all EPSCs recorded from the VS neuron (top, black) or divided into three clusters based on EPSC amplitudes (bottom, colored). A refractory period around 0 ms only occurs for EPSCs within each cluster, but not across all EPSCs. D. Schematic of three different otolith afferents converging onto one VS neuron, each eliciting EPSCs with a distinct amplitude (represented by different synaptic sizes). Right, spike activities of three afferents inferred from B.

To test this interpretation of electrophysiological data with a completely independent approach, we reconstructed the whole volume of myelinated utricular inputs onto 11 VS neurons from a high resolution re-imaged serial section EM dataset acquired from the right side of one 5.5 dpf larval zebrafish (Fig. 4A, B). We found that the connection between myelinated utricular afferents and VS neurons was relatively sparse. All VS neurons were contacted by at least two utricular afferents, but some afferents did not innervate any VS neurons (Fig. 4C). These reconstructions showed that a range of 2-6 afferents (mean±std: 3.4±1.4) converged onto each VS neuron (Fig. 4D). We compared these numbers to those derived from whole-cell physiology, where we inferred the number of convergent afferents from the number of EPSC clusters. Across all VS neuron recordings, we found a range of 0-5 afferents (1.7±1.3) converged onto each VS neuron (Fig. 4E). The result from anatomical reconstruction is largely consistent with the overall distribution of afferent contacts as measured by whole-cell physiology, presumably with some small-amplitude EPSCs elicited by the afferents not successfully clustered. Therefore, these results demonstrate that synaptic inputs from individual vestibular afferents can be separated by their stereotypic EPSC waveforms, yielding inferred afferent convergence consistent with high-resolution anatomical connectivity.

**Figure 4:**
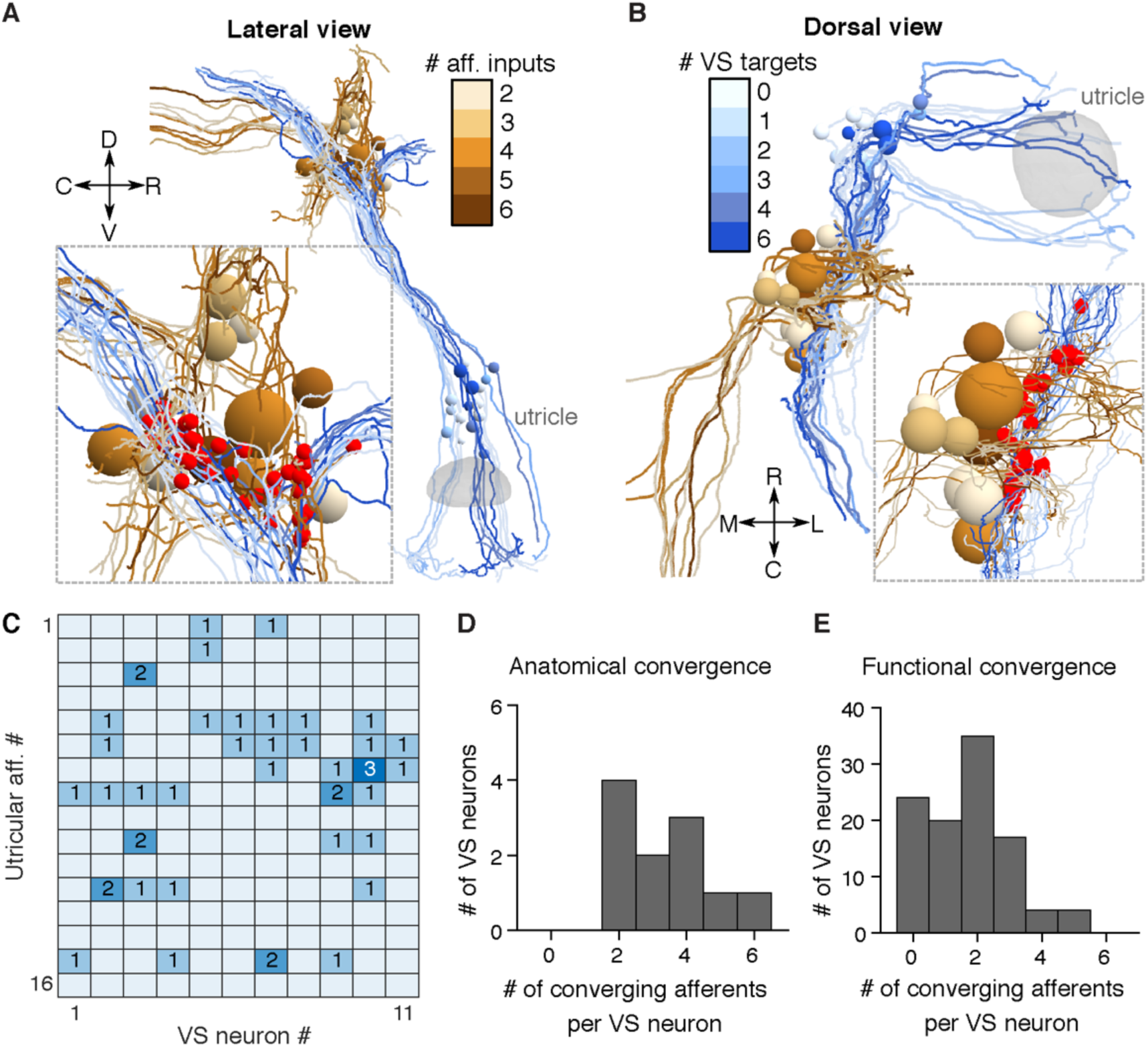
Anatomical reconstruction reveal similar convergence pattern as physiology. A. Serial-section EM reconstruction (lateral view) of all myelinated utricular afferents (blues) and VS neurons (browns) on the right side of one animal (5.5 dpf). Inset, identified synaptic contacts between afferents and VS neurons (red). Color scale represents number of distinct afferents synapsing with a given VS neuron (browns). VS neurons with greater number of afferent inputs are located more ventrally. B. Dorsal view of the same reconstruction as A. Color scale represents number of VS neurons contacted by a given afferent (blues). C. Number of distinct synaptic contacts from each utricular afferent onto each VS neuron, based on serial-section EM reconstruction. D. Histogram of the numbers of distinct afferents converging onto each VS neuron, as measured by serial-section EM reconstruction (11 neurons, 1 fish) E. Histogram of the numbers of distinct afferents converging onto each VS neuron, as inferred from whole-cell physiology recording (104 neurons, 89 fish)

### Spatial tuning of inferred otolith afferents

By recording from one VS neuron, we can infer the activity of its presynaptic afferents. This approach thus offers a unique opportunity to measure the sensory tuning of several convergent afferents simultaneously. To determine the spatial tuning of convergent afferent inputs, we delivered 2 Hz, ±0.02 g sinusoidal translational stimuli on four axes in the horizontal plane and recorded the sensory-evoked EPSCs, as shown for an example VS neuron (Fig. 5A). In this example neuron, the inferred utricular afferent (EPSC cluster) with the largest synaptic amplitude responded best to caudally-directed acceleration, while two others responded with varying sensitivities to rostrally-directed acceleration, in all cases with phase leads relative to peak acceleration (Fig. 5B). With these measurements, we can derive the preferred tuning direction, gain and phase of each afferent, as represented by the direction and length of a vector (Fig. 5B, right). To validate the consistency of the vectorial representation, we used a previously established approach (Schor et al., 1984) to quantify the tuning vectors with separately measured responses to two circular stimuli (Fig. S4 B), which showed similar preferred directions as those measured by translational stimuli (Figs. S4 A-C). Across all recordings with the animal oriented side-up, tuning of afferents was strong in the rostral (30/69) and caudal (31/69) directions, but relatively weak in the dorsal (4/69) and ventral (4/69) directions, as represented by an overlay of all inferred afferent tuning vectors (Fig. 5C). When fish were oriented dorsal-up, the axes tested were rostral-caudal and ipsilateral-contralateral (motion along an axis from one ear to the other). In this position, most afferents were strongly tuned to acceleration towards the contralateral direction (31/60), some exhibited preferential tuning to the acceleration to the rostral (4/60) and caudal (20/60) directions, and only 5/60 afferents were tuned to the ipsilateral direction (Fig. 5D). These results showed that each afferent in the larval zebrafish exhibits selective responses to different translational stimuli. Afferents overall responded best to acceleration towards the contralateral, rostral and caudal directions, which correspond to ipsilateral, nose-up and nose-down tilts in postural change (Angelaki and Cullen, 2008), consistent with the distribution of hair cell polarity in the utricular macula (Haddon et al., 1999).

**Figure 5:**
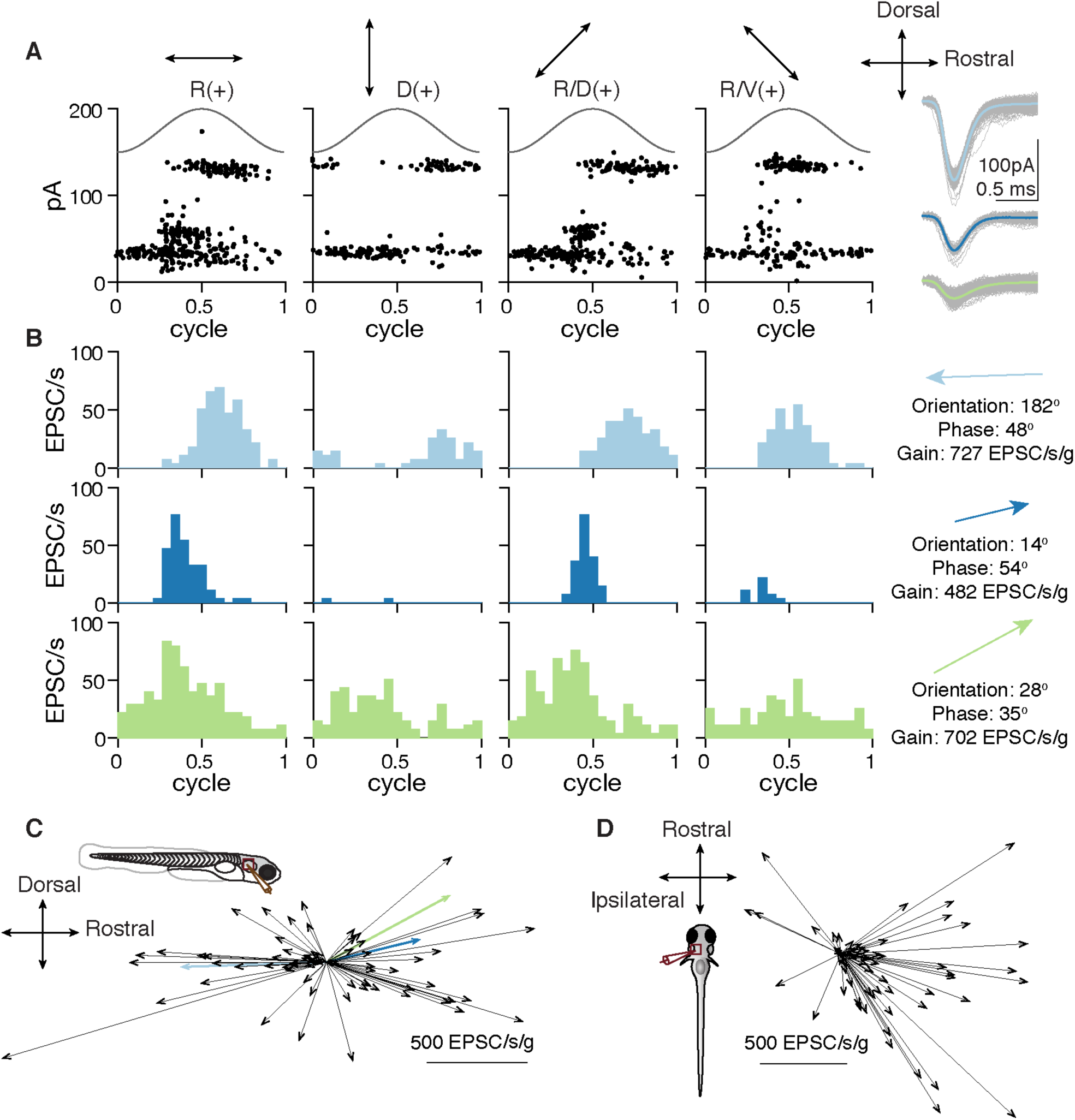
Spatial tuning of inferred otolith afferents. A. EPSC responses of an example neuron in response to 2 Hz, 0.02 g translational stimuli (solid sinusoidal line, acceleration) on 4 different axes (top, arrows). Each dot represents one EPSC; note three EPSC clusters with distinct amplitudes. Right, overlay of individual EPSCs (gray) and average (colored) for each cluster. B. EPSC tuning of three clusters. Right, vectors representing the maximum tuning direction, phase, and gain of each inferred afferent corresponding to an EPSC cluster. C. Maximum tuning directions of all afferents from VS neurons recorded from fish oriented side-up. Colored arrows represent tuning of afferents in B (69 afferents, 43 neurons, 33 fish) D. Maximum tuning directions of all inferred afferents from VS neurons recorded from fish oriented dorsal-up. (60 afferents, 36 neurons, 36 fish)

### Temporal tuning of inferred otolith afferents

The sensitivity and phase of vestibular afferents varies for motion at different frequencies (Fernandez and Goldberg, 1976b). The tuning of otolith afferents ranges from typically more jerk-encoding (derivative of acceleration) at low frequencies to more acceleration-encoding at high frequencies. What temporal tuning profile do afferents in larval zebrafish exhibit? We applied translational stimuli with different frequencies (0.5-8 Hz, ±0.02 g) on the rostral-caudal axis. In the example neuron, all three inferred otolith afferents showed similar tuning, with progressively stronger responses with increasing frequencies of stimulation (Fig. 6A). Across group data acquired at both ±0.02 g and ±0.06 g, the average tuning gain increased 3.3-fold (0.02 g) and 2.3-fold (0.06 g) from 0.5 Hz to 8 Hz (Fig. 6B). Most afferents (39/48) showed at least 2-fold increase from 0.5 Hz to 4 Hz in tuning gain at either 0.02 g or 0.06 g. Only one afferent had relatively flat gain (< 50% increase) at both 0.02 g and 0.06 g, and its tuning was overall weak (mean gain: 1.88 and 2.24 EPSC/s respectively), suggesting it was less sensitive or not tuned on the rostral-caudal axis. Regardless of tuning direction (rostral: 44%, 21/48; caudal: 56%, 27/48), afferents exhibited a phase lead relative to peak acceleration at various tested stimulus magnitudes and frequencies ((Fig. 6C, S5). On average, the phase lead at low frequency (0.5 Hz) was 84.0° for 0.02 g and 78.6° for 0.06 g. At high frequency (8 Hz), the phase lead was reduced to 33.6° for 0.02 g and 39.3° for 0.06 g. The temporal dynamics of the afferents resembled those of previously reported irregular units (Goldberg et al., 1990), with low spontaneous firing rates (10.28±9.1 EPSC/s) and larger coefficients of variation (CV). The average CV across inferred afferents was 0.97±0.24, and the smallest CV was 0.5 (Fig. S6), indicating that no regular-firing otolith afferents were detected synapsing onto VS neurons. We conclude that the otolith afferents act as a high-pass filter, encoding a mixture of acceleration and jerk, similar to otolith afferents in primates (Laurens et al., 2017).

**Figure 6:**
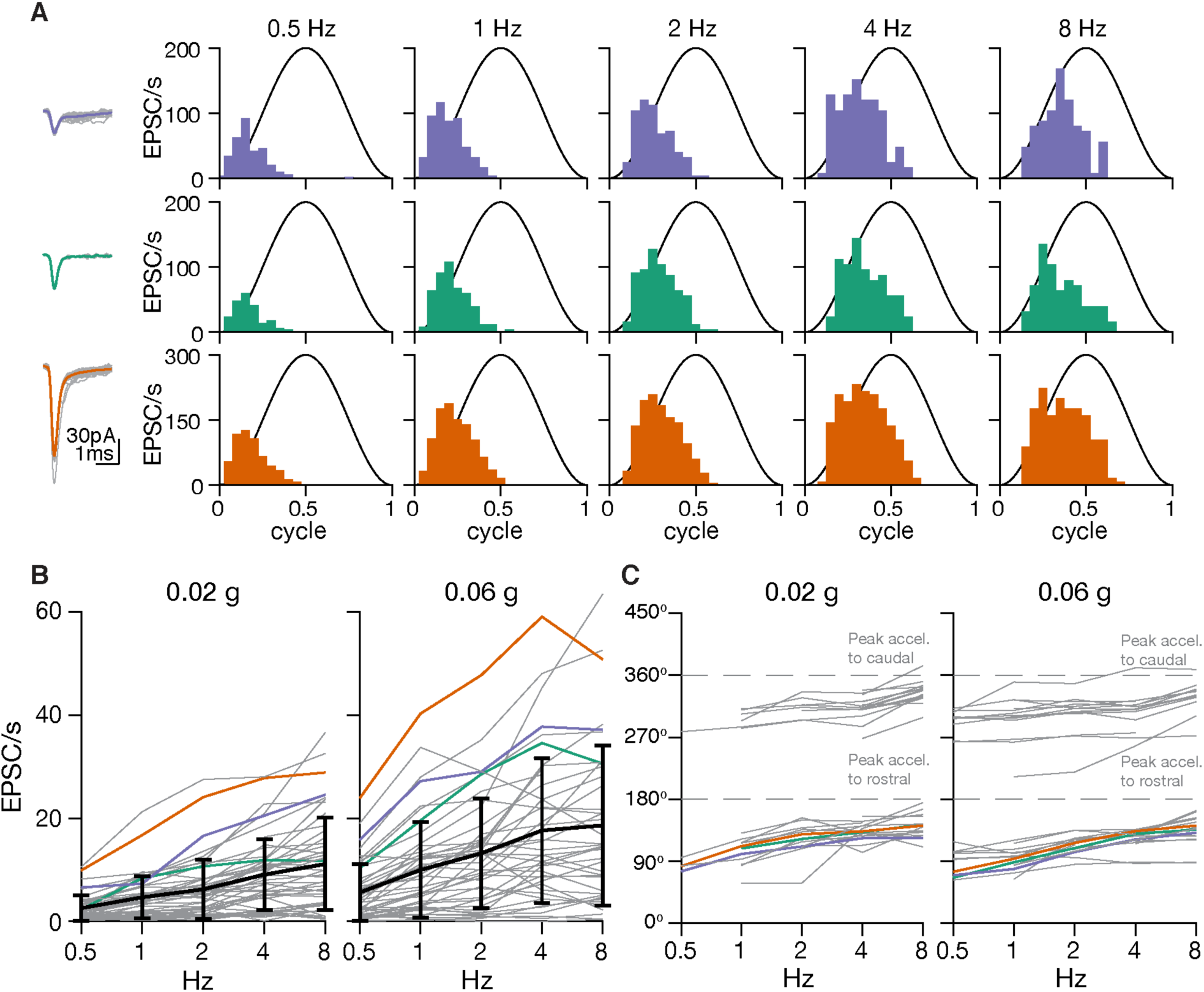
Temporal tuning of inferred otolith afferents. A. Sensory tuning of afferent inputs to one VS neuron during translational movement at 5 different frequencies in the rostral(+)-caudal(-) axis. Left, EPSC waveforms of three different clusters recorded from one VS neuron. Right, temporal tuning profile of each EPSC cluster on the rostral-caudal axis. B. Gains of inferred afferents across different frequencies of translational acceleration. Gray, individual afferents; colored, afferents from A; black, mean and standard deviation of gains from all afferents (0.02 g, 48 afferents; 0.06 g: 46 afferents; 25 neurons, 20 fish) C. Phases of inferred afferents across frequencies, relative to sinusoidal stimulus. 180° (0.5 cycle in A) represents the peak of acceleration towards rostral direction; 360° represents the peak of acceleration towards caudal direction (0 or 1 cycle in A). Data were thresholded to only include afferents whose gain was > 5 EPSC/s (0.02 g, 36 afferents; 0.06 g, 38 afferents; 25 neurons, 20 fish)

### Preferential convergence

Individual VS neurons can receive inputs from afferents with similar (Fig. 6A) or different tuning (Fig. 5B). Is afferent tuning convergence random or structured? The responses of inferred afferents that converge onto the same VS neuron were represented by their tuning vectors (Fig. 7A). The angle between the vectors indicates the similarity of convergent inputs, with a small angle for a VS neuron with similarly tuned inputs and a large angle for a VS neuron with differently tuned inputs. From 43 VS neurons recorded in the side-up orientation, 60% (38/63) of converging afferent pairs had small angles (<45°) and 27% (17/63) had large angles (>135°). Compared to a random pairing angle distribution generated by bootstrapping, the percentage of similarly tuned convergent afferent pairs was significantly higher than chance (Fig. 7B, left). From 36 VS neurons recorded in the dorsal-up orientation, there were 71% (37/52) of inferred afferent pairs with a converging angle smaller than 45°, and only 2% (1/52) with a converging angle larger than 135° due to the small number of ipsilaterally tuned afferents (Fig. 5D). Nonetheless, the probability of similarly tuned afferent convergence (<45°) was significantly higher than that chance (Fig. 7B, right). For afferent pairs with converging angle larger than 45° (45°-90°, 90°-135°, 135°-180°), their probabilities was slightly lower than their respective estimated distribution by bootstrapping. Accordingly, on a given body axis (R-C or I-C), convergent afferents are also more likely to encode similar tuning directions (Fig. 7C). These results suggest that afferents with similar tuning direction preferentially converge at rates exceeding what would be expected by random connectivity.

**Figure 7:**
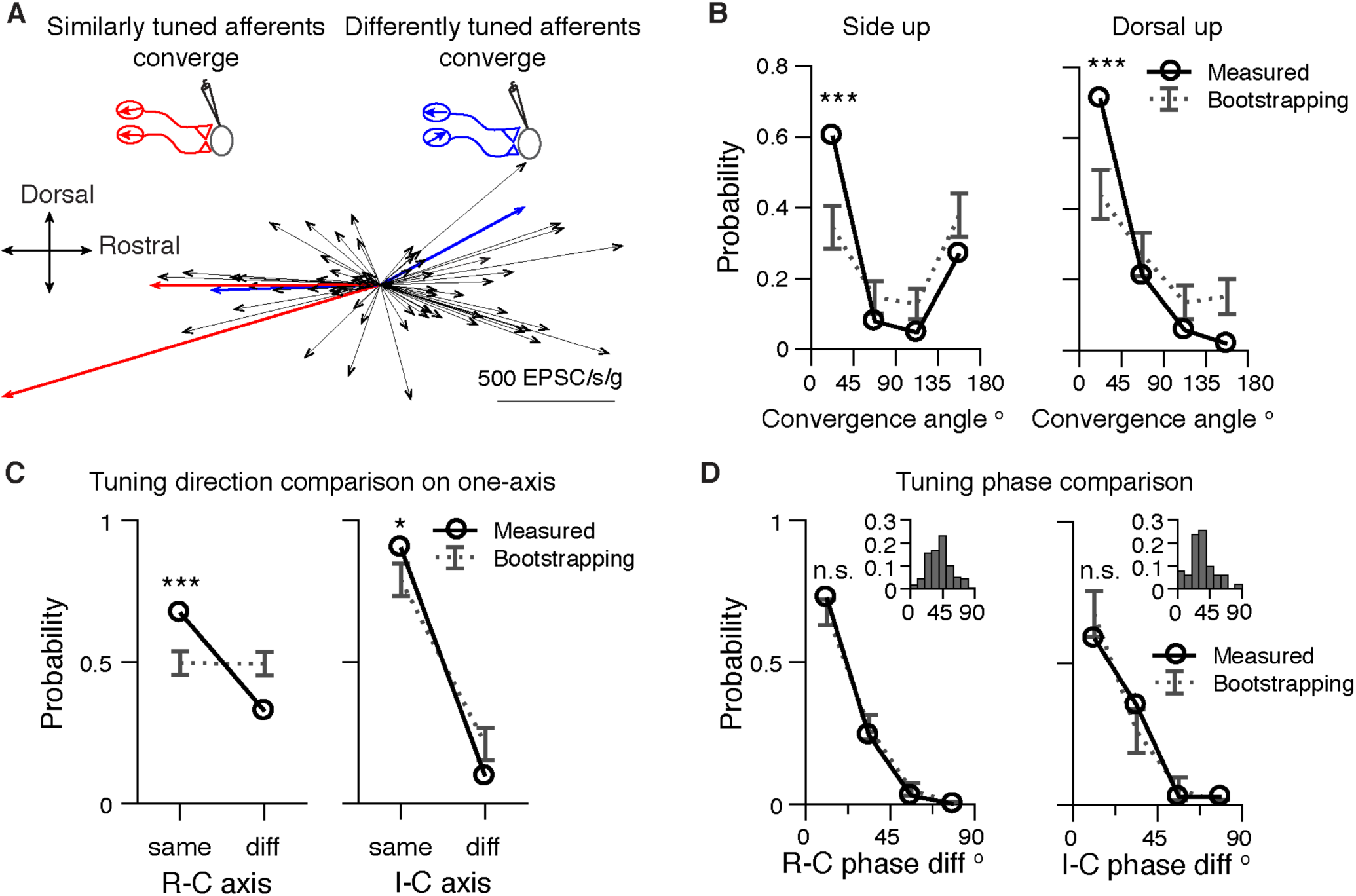
Afferents with similar tuning direction preferentially converge. A. Example of two pairs of converging afferents from two VS neurons in side-up fish. Red, converging afferents are similarly tuned, with small convergent angle between the pair; Blue: converging afferents are differently tuned, with large convergent angle between the pair. B. Probability distribution of converging angles for measured and randomly generated afferents pairs in side up fish and dorsal up fish. Two tailed z-test, side up, 0-45°: p=2e-5, 135-180°: p=0.08. (63 afferent pairs); dorsal up, 0-45°: p=8e-5, 135-180°: p=0.007. (52 afferent pairs) C. Probability distribution of converging afferents tuned to the same direction vs different direction, on the rostral-caudal and ipsilateral-contralateral axis. Two tailed z-test, R-C, same: p=1e-5, diff: p=4e-5 (150 afferent pairs); I-C, same: p=0.044, diff: p=0.044 (52 afferent pairs). D. Probability distribution of phase difference for converging afferents, on the rostral-caudal and ipsilateral-contralateral axis. Two tailed z-test, R-C, 0-22.5° p=0.26 (103 afferent pairs); I-C, 0-22.5°, p=0.28 (34 afferent pairs). Inset: distribution of tuning phase of afferents, R-C, 177 afferents; I-C, 60 afferents; 90° represents the peak of acceleration of preferred direction (2 Hz, 0.02 g).

Do converging afferents also have similar tuning phase regardless of their tuning direction? Most afferents are phase-leading with 2 Hz, 0.02 g stimulation (Figs. 6 and 7D, inset), and the phase difference between afferents is small (R-C: 41° ±16°, n=177, I-C: 33° ±17°, n=60). Consequently, most afferent pairs (R-C, 68±4.6%; I-C, 68±8%) selected randomly have very small phase difference (phase diff. < 22.5°) (Fig. 7D). Both the probability of converging afferents having similar phase (phase diff. < 22.5°) (R-C, 73%, 75/103; I-C, 59%, 20/34) and the cumulative distribution (Fig. S7) lay within the bootstrap predications on the rostral-caudal and ipsilateral-contralateral axes. Therefore, tuning phase between converging afferent pairs is similar, in accordance with their relatively homogeneous distribution.

In conclusion, we found that afferents forming synaptic connections with the same postsynaptic VS partner typically have similar spatiotemporal tuning properties. In particular, afferents with similar tuning direction preferentially converge, which explains the long-standing observation that most VS neurons exhibit simple cosine tuning (Peterson, 1970, Schor et al., 1984). However, a non-negligible number of VS neurons receive convergent input from differently tuned afferents, a potential source for the complex spatiotemporal tuning of central vestibular neurons.

### Complexity of central tuning is determined by the similarity of afferent inputs

Complex sensory tuning of central neurons is thought to arise from convergence of more simply tuned inputs with differing spatial and temporal properties, in vestibular (Angelaki et al., 1993), as well as visual (Jia et al., 2010) and somatosensory (Petersen, 2007, Roy et al., 2011) systems. For example, complex tuning such as bidirectional (Peterson, 1970) and broadly tuned sensory responses (Angelaki, 1992) of central vestibular neurons can be computationally reconstructed from multiple modelled cosine-tuned inputs. However, directly measuring these inputs has been technically difficult, and it is unclear whether such models can sufficiently explain the activity of central neurons. Therefore, we took advantage of the inferred afferent spiking to examine whether the tuning of VS neurons can be constructed from the convergence of otolith afferents.

We observed that different VS neurons showed simple or complex membrane potential responses to translational stimuli on the rostral-caudal axis. An example simple cell was only depolarized during a specific phase of acceleration (Fig. 8A), whereas a complex cell exhibited multiple depolarized periods during the stimulus (Fig. 8C). Next, we measured the EPSC tuning in the same VS neurons. In the example simple neuron, sensory evoked EPSCs exhibit three distinct amplitudes (Fig. 8B), indicating three afferents converge onto the cell. These three afferents showed similar tuning to each other, with strongest responses for rostrally-directed acceleration. In contrast, the four inferred afferents that converge onto the example complex cell exhibited a different tuning pattern. Two afferents were tuned to rostrally-directed acceleration and the other two to caudally-directed acceleration (Fig. 8D).

**Figure 8:**
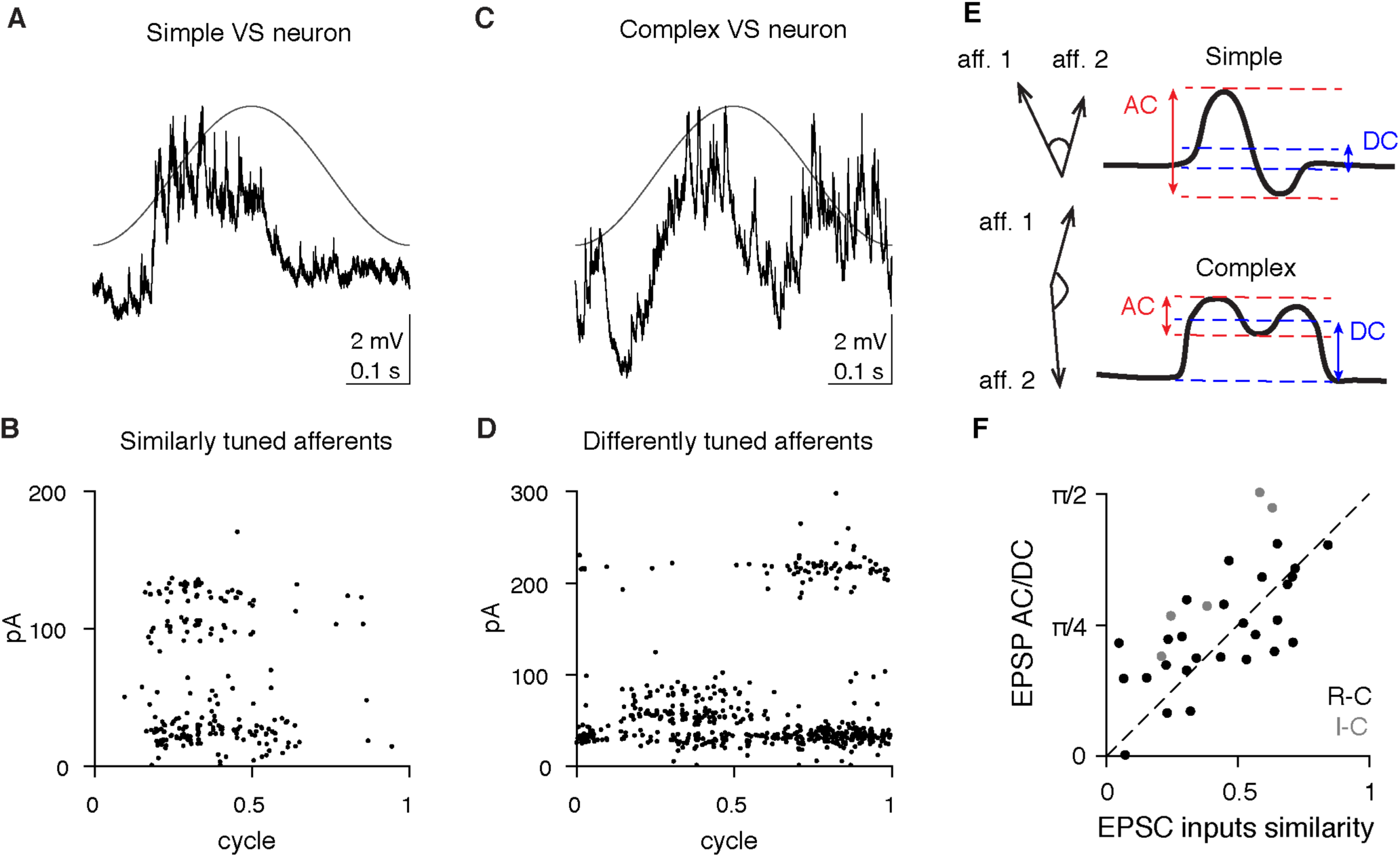
Subthreshold tuning responses of VS neurons are explained by the similarity of tuning of afferent inputs. A. Average membrane potential change in a VS neuron with simple response to 2 Hz, 0.02 g translational movement on the rostral(+)-caudal(-) axis. B. EPSC responses for the simple cell shown in A. All inferred afferents exhibit similar tuning. C. Average membrane potential change in a VS neuron with complex subthreshold response to 2 Hz, 0.02 g translational movement on the rostral(+)-caudal(-) axis D. EPSC responses for the complex cell shown in B. Different inferred afferents exhibit distinct tuning, as shown by the different temporal responses of the large amplitude and medium amplitude EPSCs. E. Schematic of different magnitudes of AC and DC responses in a simple and complex cell. Left, the hypothesized models of similarly tuned or differently tuned convergent afferent inputs underlying simple or complex responses, respectively. F. Correlation of EPSC inputs similarity index and EPSP AC/DC response ratio (see Methods), for all non-spiking VS neurons with multiple convergent afferents. Sensory tuning of afferent inputs and EPSPs was measured on the R-C axis (black, n=27) and I-C axis (grey, n=5). Dashed, unity line. Pearson’s R: 0.67

To examine this relationship across the population, we defined an afferent inputs similarity index for multiply innervated VS neurons, based on the phase of afferent inputs and their EPSC amplitudes. The index ranges from 0-1, with smaller index representing more divergent ESPC input tuning and larger index representing more similar tuning (see Methods). A classifier originally developed for visual cortical neurons was used to quantify the tuning complexity of the postsynaptic neuron’s membrane potential responses to sensory stimuli (Skottun et al., 1991). In this metric, neurons with simple tuning show large AC and small DC responses, whereas complex cells exhibit small AC and large DC responses (Fig. 8E). We found that the AC/DC ratio of the membrane potential was strongly correlated with the similarity index of afferent inputs (Fig. 8F). In other words, convergence of more similarly tuned afferents yields a more simple VS neuron response, and the convergence of more differently tuned afferents generates a more complex postsynaptic response.

### Spike tuning generation from similar and differently tuned afferents

We next extended the comparison of presynaptic to postsynaptic tuning by measuring the spiking responses of VS neurons during sensory stimulation. In a subset of VS neurons, the largest translational stimuli that we could deliver while holding the cell was sufficient to evoke postsynaptic firing; in other neurons, a small bias current was injected to evoke spiking during sensory stimulation (see Methods). Most VS neurons exhibited simple spike tuning, and received convergent inputs from similarly tuned afferents (Fig. 9A). Some VS neurons with simple spike responses received convergent inputs from differently tuned afferents (Fig. 9B). Finally, complex spike tuning in VS neurons was always generated by inputs from differently tuned afferents (Fig. 9C). These three categories of input-output transformation (similar to simple, different to simple, different to complex) were identified across all recordings from VS neurons (Fig. 9D). In total, most recordings (R-C, 6/13; I-C, 14/19) exhibited simple spike tuning, of which 69% (20/29) received inputs from similarly tuned afferents, and 31% (9/29) received inputs from differently tuned afferents (Fig. 9E). Thus, convergence of similarly tuned afferents yields simple spike tuning, but convergence of differently tuned afferents can yield either simple or complex spike tuning. Interestingly, complex spike tuning was only observed on the R-C axis (3/13 recordings), which subserves pitch movements, not the I-C axis (0/19), which subserves roll (Fig. 9F). These results indicate that VS neurons may play different roles in maintaining body balance on different body axes.

**Figure 9:**
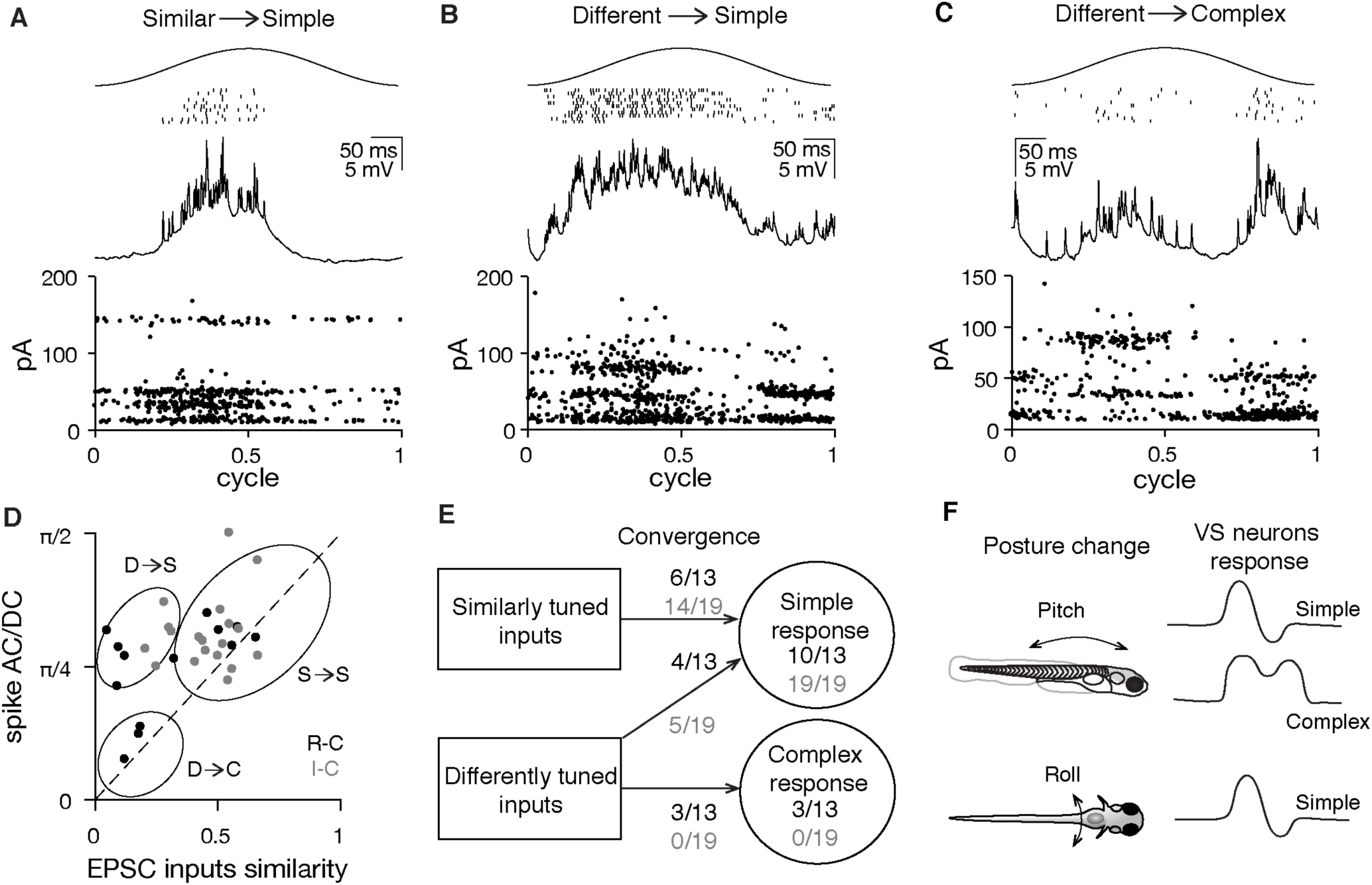
Spiking tuning complexity of VS neuron is partially explained by the similarity of tuning for the afferent inputs. A. Example cell showing that simple spiking tuning response is constructed from afferent inputs with similar tuning direction. Top, sensory-evoked spike raster of a VS neuron during 2 Hz, 0.02 g translational movement on the I(-)-C(+) axis (11 trials); middle, average membrane potential of the VS neuron (11 trials); bottom, sensory-evoked EPSC response (12 trials); each dot represents one EPSC. B. As in A, for an example cell showing simple spiking tuning response arising from afferent inputs with different tuning directions on the I(-)-C(+) axis. C. As in A, for an example cell with a complex spiking tuning response due to afferent inputs with different tuning directions on the R(+)-C(-) axis. D. Correlation of EPSC input similarity index and spike activity AC/DC ratio, for all spiking VS neurons on the R-C axis (black, n=13 recordings) and I-C axis (gray, n=19). Circles, three categories of input-output transformation, corresponding to examples in A, B and C. Dashed, unity line. Pearson’s R: 0.51 E. Quantification of input-output transformation of VS neurons; Ns represent individual cells. F. Summary of different responses of VS neurons responding to posture change on the pitch and roll axes.

## Discussion

### Sensory convergence in the central vestibular nuclei

Taking advantage of the invariant synaptic transmission of electrical synapses, we separated distinct afferent inputs that converge onto VS neurons and measured the spatial and temporal tuning of each converging afferent *in vivo*. This analysis is facilitated by the sparseness of connectivity, with < 6 afferents synapsing with each VS neuron. These data resolve a conflict in the literature: anatomically, very few otolith afferent terminals are observed in the lateral vestibular nucleus (Newlands and Perachio, 2003), but physiologically, afferent stimulation elicits monosynaptic EPSPs in VS neurons (Boyle et al., 1992). Our data reveal that sparse but powerful afferent synaptic contacts, located on the lateral dendrites of VS neurons, are sufficient to drive the membrane potential of the cell during sensory stimulation. Although these large afferent inputs drive sensory responses, VS neurons receive a wealth of non-vestibular synaptic contacts on their large dendritic arbors. This is consistent with previous findings that the activity of VS neurons is regulated by locomotion (Orlovsky, 1972), proprioception (Neuhuber and Zenker, 1989), and other inputs (Sarkisian, 2000, Witts and Murray, 2019). Interestingly, lateral geniculate neurons (LGN) of the visual thalamus display a similar pattern of connectivity, with sparse, powerful afferent inputs from retinal ganglion cells and weaker, diverse inputs from other sources (Usrey et al., 1999, Sherman, 2005). Our findings suggest that this configuration is common to VS neurons as well.

Similarly tuned otolith afferents preferentially converge onto VS neurons (Fig. 7), demonstrating that feedforward excitation can generate central neurons with simple response properties. In a similar vein, thalamocortical inputs with similar angular tuning also preferentially project onto the same site in somatosensory cortex, and the preferred tuning direction of the cortical neuron can be predicted by that of the presynaptic thalamic neuron (Bruno et al., 2003). Furthermore, we found that convergence of differently tuned afferents can yield a more complex postsynaptic response in central vestibular neurons, similar to bidirectional or complex tuning observed previously in cats (Peterson, 1970) and primates (Angelaki and Dickman, 2000). This result generally supports the hypothesized model (Angelaki, 1992) that the tuning of central vestibular neurons can be constructed from cosine tuned inputs with varying tuning properties. However, we find that convergence of differently tuned afferents can also yield simple tuning in VS neurons (Fig. 8B), suggesting other factors such as inhibition (Straka and Dieringer, 1996) and thresholding (Priebe et al., 2004) might be involved. We found no evidence for polysynaptic excitatory circuits during afferent stimulation (Fig. S9 A and B), and modelling indicates that excitatory synaptic input is sufficient to predict subthreshold membrane potential and tuning (Fig. S9 C-F). However, stronger stimuli might elicit inhibition and other nonlinearities. Across brain regions, sensory tuning of central neurons is constructed by a variety of mechanisms. These include afferent convergence pattern (Alonso and Martinez, 1998, Priebe and Ferster, 2012), local excitatory or inhibitory modulation (Wilent and Contreras, 2005), and nonlinear dendritic computation (Lavzin et al., 2012). Our results demonstrate that sensory response of a central neuron can be constructed from the afferent inputs in a direct feedforward manner.

### Otolith afferent tuning properties in the larval zebrafish

The derived spatial tuning profile of afferents in the larval zebrafish is similar to the polarity of the hair cells in otolith macula, consistent with results in fish (Fay, 1984, Platt, 1977) and primates (Fernandez et al., 1972). Notably, tuning to dorsal or ventral acceleration was relatively weak for most afferents, presumably due to the horizontal orientation of the utricular membrane in larval zebrafish inner ear. Afferents were preferentially tuned to contralateral acceleration (ipsilateral tilt) in the roll axis, consistent with the dearth of ipsilaterally tuned hair cells in larval zebrafish (Haddon et al., 1999). In species with more centrally located line of polarity reversal (Fernandez and Goldberg, 1976a, Tomko et al., 1981), we would predict more convergence of oppositely tuned afferents, and correspondingly more complex response of VS neurons in the roll axis, as seen in cats (Peterson, 1970).

A significant question in vestibular systems is whether central vestibular neurons receive selective projections from afferents with regular as opposed to irregular firing. Both regular and irregular afferents are thought to converge on VS and vestibulo-ocular reflex neurons in mammals, based on studies comparing recruitment thresholds of afferent inputs (Boyle et al., 1992). Our data provide direct evidence that vestibular inputs to VS neurons exhibit classic characteristics of irregular afferents (Eatock and Songer, 2011): high-pass tuning, low spontaneous firing rate, and high CV of firing (Fig. S6). It is unknown whether regular utricular afferents exist in the larval zebrafish. Although regular utricular afferents were observed in guitarfish (Budelli and Macadar, 1979), they appear absent in toadfish (Maruska and Mensinger, 2015) and sleeper goby (Lu et al., 2004). Based on serial section EM, many afferents make no contacts with VS neurons (Fig. 4C), leading us to conclude that either regular afferents have not yet developed or that they do not contact VS neurons in the larval zebrafish.

### Linear and fast synaptic transmission via gap junctions

Our data reveal that electrical synapses mediate linear synaptic transmission from otolith afferents to the VS neurons. In contrast, synaptic transmission at retinotectal afferents in larval zebrafish is mediated solely by glutamate (Smear et al., 2007), suggesting that electrical synapses in vestibular afferents are perhaps not simply a feature of early larval development but play an important role in circuit computations, potentially via their amplitude invariant transmission. Interestingly, mammalian vestibular afferent synapses also exhibit amplitude invariant transmission in the medial vestibular nucleus (Bagnall et al., 2008) and cerebellum (Arenz et al., 2008, Chabrol et al., 2015), but via specialized glutamatergic terminals (Turecek et al., 2017, McElvain et al., 2015), indicating that frequency-independent transmission is a hallmark of vestibular signaling across vertebrates. Furthermore, mixed electrical and chemical synapses have been anatomically identified between vestibular afferents and VS neurons in both adult fish (Korn et al., 1977) and rodents (Nagy et al., 2013), suggesting the mixed synapse may be a conserved mechanism across species to implement fast, frequency-independent transmission in the lateral vestibular nucleus. The amplitude invariance of this connection allowed us to examine whether there was any relationship between an afferent’s sensory gain or firing rate and the synaptic amplitude it evokes in a VS neuron. No correlation appeared in either of these measures (Fig. S8), indicating that at least within this population, synapse size is not “normalized” by firing rate.

### VS pathway underlying sensorimotor transformation

The VS pathway is important for posture control. Larval zebrafish swim at high frequencies up to 100 Hz (McLean and Fetcho, 2009) and are naturally unstable in water (Ehrlich and Schoppik, 2017). Our study examined the response of VS neurons with translational stimuli in the range of 0.04-0.12 g and 0.5-8 Hz, head movement parameters comparable to slow swimming (Voesenek et al., 2016) or small angle tilting motion in the larval zebrafish. In the roll axis all VS neurons are tuned to ipsilateral tilt (Fig. S10), consistent with data from calcium imaging (Migault et al., 2018, Favre-Bulle et al., 2018), suggesting they might excite specific motor units in the spinal cord to produce compensatory movements (Bagnall and McLean, 2014). On the pitch axis, VS neurons have more heterogeneous responses, including simple tuning to either rostral or caudal acceleration (Fig. S10), as well as complex responses (Fig. 9E) to both directions. Thus, when the animal is destabilized by excessive nose-up or nose-down tilts, VS neurons might activate non-specific motor units, increasing the likelihood of swim bouts to regain balance (Ehrlich and Schoppik, 2017).

The high-pass tuning and phase lead of otolith afferents innervating VS neurons will make larvae most sensitive to ongoing changes in tilt or acceleration, especially at high frequency. These data are consistent with behavioral observations that larvae become more likely to swim to correct their position in the pitch axis when angular velocity (i.e., changing tilt) reaches a critical threshold (Ehrlich and Schoppik, 2017). This compensatory postural adjustment, which relies on both trunk and fins, is absent in *rock solo* larvae (Ehrlich and Schoppik, 2019), in line with our results on the absence of sensory tuning in those animals. Larval VS neurons receive similar amounts of inputs from rostrally and caudally tuned afferents, suggesting both nose-up and nose-down tilts are equally detected by the VS pathway. In contrast, the vestibulo-ocular pathway shows an anatomical bias for representation of nose-up body tilt (Schoppik et al., 2017). This indicates that different strategies might be involved to adjust body posture and eye position for pitch movements in larval zebrafish.

Moreover, it is important for animals to distinguish self-generated and external vestibular signals. We described the direct excitatory inputs from vestibular afferents onto the VS neurons during passive movements. How do self-generated motion signals modulate the activities of VS neurons? Projections from Purkinje cells in the cerebellum are thought to suppress sensory-evoked activity in VS neurons during voluntary self-motion (Cullen, 2019). In the future, it would be interesting to use *in vivo* whole-cell physiology to investigate how central vestibular neurons distinguish self-generated movements from passive movements.

## Methods

### Fish lines and husbandry

*Tg(nefma:LRL:Gal4)* was established by injecting the construct containing hsp70 promoter (Kimura et al., 2014), and the insertion site was set at the upstream of the nefma gene with the CRISPR target sequence: CATCGACGGATCAATGG. The *Tg(nefma:gal4)* fish line was generated by crossing *Tg(nefma:LRL:Gal4)* with a ubiquitous-Cre fish. The *otogc.1522+2T>A -/-* (rock solo) mutant is vestibular deficient due to a splice site mutation in *otogelin* (Mo et al., 2010, Roberts et al., 2017). Rock solo homozygotes on a *Tg(nefma:gal4, UAS:GFP)* background were crossed to rock solo WT/heterozygotes to produce clutches containing WT, heterozygotes and homozygotes for recording purposes. The rock solo homozygotes were identified by the absence of anterior otolith (utricle). All experiments and procedures were approved by the Animal Studies Committee at Washington University and adhere to NIH guidelines.

Animals were raised and maintained in the Washington University Zebrafish Facility at 28.5°C with a 14:10 light:dark cycle. Larval zebrafish (4-7 dpf) were housed either in petri dishes or shallow tank with system water. Adult animals were maintained up to 1 year old with standard procedure.

### Electrophysiology

VS neurons were targeted for whole-cell patch clamp recordings based on their characteristic position and fluorescence in the *Tg(nefma:gal4, UAS:GFP)* fish. The larvae (4-7 dpf) were paralyzed by 0.1% *α*-bungarotoxin and embedded in a 10 mm FluoroDish (WPI) with low-melting point agarose (Camplex SeaPlaque Agarose, 2.4% in system water). Fish were immersed in extracellular solution ([in mM] NaCl 134, KCl 2.9, MgCl_2_ 1.2, HEPES 10, glucose 10, CaCl_2_ 2.1, osmolarity ∼295 mOsm and pH ∼ 7.5) and a small piece of skin above the brainstem was carefully removed by sharpened tungsten pins. The fish was transferred to an epifluorescence microscope equipped with immersion objectives (Olympus, 40x, 0.8 NA), infrared differential interference contrast optics and air-bearing sled recording table.

Patch pipettes (7-9 MΩ) were filled with internal solution ([in mM] K gluconate 125, MgCl_2_ 2.5, HEPES 10, EGTA 10, Na_2_ATP 4, Alexa Fluor 568 or 647 hydrazide 0.05-0.1, osmolarity ∼295 mOsm and pH ∼ 7.5). After whole-cell configuration was achieved, voltage clamp and current clamp signals were recorded at room temperature with a Multiclamp 700B, filtered at 10 kHz (current clamp) or 2 kHz (voltage clamp), digitized at 50 kHz with Digidata 1440 (Axon Instruments), and acquired by Clampex 10 (Molecular Devices).

Before the vestibular stimulus was delivered to the fish, the immersion objective was removed from the recording chamber. During the recording, series resistance was monitored every 15 s to ensure good recording quality; neurons with series resistance variation > 25% were discarded. After recording, the recorded cell was imaged with epifluorescence to confirm cell identity.

### Vestibular stimulation

The recording rig was custom-designed to allow delivering user-controlled movement to the fish during recording without losing whole-cell access. The microscope and a one-axis or two-axis air-bearing sled (Aerotech, ABL1500WB or ABL1500&1500WB) were fixed on an air table. Manipulators (Microstar) and recording platform (ThorLab) were positioned on the sled. The sled was powered with the Aerotech transformers (TM5), NPdrivers (NDRIVECP10-MXU&NDRIVECP20-MXU) and nitrogen gas (Airgas, NI UHP300). Stimuli were designed in Matlab and imported into the program by Aerotech software (Motion Designer/Composer), with additional tuning as required to compensate for the motion of the underlying air table. Movement was recorded by an accelerometer (Sparkfun, ADXL335) attached to the platform. Motion signals were digitized at 50 kHz with Digidata 1440 (Axon Instruments), and acquired in Clampex 10 (Molecular Devices).

Fish were embedded either dorsal side up (movements on rostral-caudal and ipsilateral-contralateral axes) or left/right side up (movements on dorsal-ventral and rostral-caudal axes) and a series of frequency-varying sinusoidal translational stimulus was applied. The stimulus amplitude was set at 0.02 g or 0.06 g (min to max: 0.04 g or 0.12 g respectively), and stimulus frequency range was 0.5-8 Hz. For spatial tuning measurements, linear translation was applied on four different axes (0-180°, 45°-225°, 90°-270°, 135°-315°) on the horizontal plane. To record spike tuning in neurons without spontaneous firing, a rheobase current was injected to depolarize the cell.

### Vestibular afferent stimulation and pharmacology

A glass pipette electrode (2-5 MΩ) filled with extracellular solution ([in mM] NaCl 134, KCl 2.9, MgCl_2_ 1.2, HEPES 10, glucose 10, CaCl_2_ 2.1, osmolarity ∼295 mOsm and pH ∼ 7.5) was connected to a stimulator (A-M systems, Model 2100), and placed in the vestibular ganglion to stimulate the vestibular afferents. A train of 0.1 ms, 1 *µ*A - 1 mA electrical pulses at varying frequencies were delivered to elicit EPSCs in the recorded cells. At least 20 trials of evoked EPSCs were recorded to establish a stable baseline. AMPA receptors and gap junctions were blocked with 10 *µ*M NBQX and 500 *µ*M carbenoxolone, respectively.

### Electron Microscopy

Ultrathin serial sections of brainstem from a 5.5 dpf zebrafish were a generous loan from J. Lichtman and F. Engert. Using the published 18.8 nm/px reference map and reconstructions (Hildebrand et al., 2017), we targeted re-imaging at 4 nm/px to the entirety of the myelinated utricular afferents (identified by their peripheral processes reaching for the utricular macula) and VS neurons (identified by their position and axonal projections into the spinal cord) on one side of the brainstem, covering ∼95 *µ*m in the rostrocaudal axis. Imaging was carried out on a Zeiss Merlin 540 FE-SEM with a solid-state backscatter detector. The ATLAS scan engine was controlled via WaferMapper (Hayworth et al., 2014). The resulting images were aligned onto the 18.8 nm/px dataset using linear affine transformations in FIJI with the TrakEM2 plug-in (Cardona et al., 2012) and will be freely available after publication. In a small subset of identified synapses, we carried out further re-imaging at 1nm/px to visualize the hallmarks of gap junctions.

The existing tracings of VS neurons and utricular afferents were extended to cover branches that had been missed or untraced in the original dataset. Afferent/VS neuron appositions were considered to be synaptic contacts if the presynapse contained vesicles, the membranes were tightly apposed and straight, and there were signs of a postsynaptic density. In cases where appositions were more difficult to determine, such as those parallel to the plane of section, vesicle clustering at a tight apposition was used as the criterion for a synapse.

### Analysis

All analysis are implemented in Matlab (Mathworks).

### Event detection

EPSC events were detected by a derivative method (Bagnall and McLean, 2014). Tuning index of all EPSCs was calculated as the vectoral sum of all events’ phase, weighted by the EPSC amplitude.

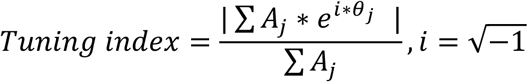

(*A*_*j*_ is the amplitude of each EPSC event *j*, and *θ*_*j*_ is the phase of that event relative to the sinusoidal stimulus on each axis.)

### Deconvolution of electrical and chemical signals

We assumed that the signals we observed on voltage clamp were majorly composed of electrical EPSCs and chemical EPSCs from afferents, based on our observation from the pharmacology data.

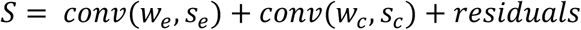

(*w*_*e*_, *w*_*c*_ are the kernels of electrical and chemical components of EPSCs, both derived from their waveforms shown in Fig. 2, and *s*_*e*_, *s*_*c*_ are the separated electrical and chemical signals)

A sparse deconvolution algorithm with L1 regularization, derived from FISTA ((Beck and Teboulle, 2009), was applied to obtain *s*_*e*_, *s*_*c*_ by minimizing the objective function:

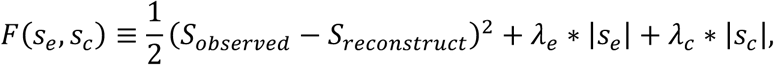

where:

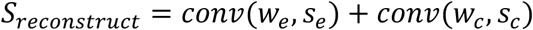

*λ*_*e*_ and *λ*_*c*_ were defined by the root mean square of the signal and magnitude of kernel waveform: 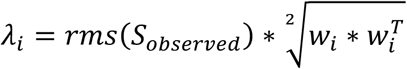. Maximum iteration cycle was set at 500

### Clustering and quantification of electrical events

Amplitude-invariant EPSCs are mediated by gap junctions, therefore only electrical signals *s*_*e*_ were used to infer individual afferent inputs. A threshold of 3.5 * *std*(*s*_*e*_) was used for event detection. Detected electrical events were clustered by ISO-SPLIT (Chung et al., 2017). Some clusters were merged or split manually after examination. Clusters with refractory period (threshold: probability < 0.003 within 1 ms) in auto-correlograms (100 ms) were considered from an individual afferent.

For each cluster, the tuning vector of inferred afferent *k* on each axis was quantified as:

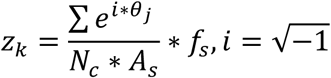

(*θ*_*j*_ is the phase of EPSC event *j* in cluster *k. N*_*c*_ is the number of cycles for sinusoidal translation, *f*_*s*_ [Hz] and *A*_*s*_ [g] are the frequency and amplitude of the stimulus. The absolute value and argument of *z* represent the tuning gain and the tuning phase, respectively.)

Tuning in four axes was fitted into a 2-dimensional spatiotemporal model (Angelaki, 1992) to obtain the maximum tuning direction, the tuning gain and phase in that direction.

Afferent inputs similarity index for a VS neuron was determined as:

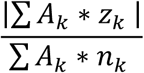

(*A*_*k*_, *z*_*k*_, *n*_*k*_ are the average EPSC amplitude, tuning vector and number of events for cluster *k*.)

### AC/DC response quantification

AC of membrane potential and spiking response were defined as the amplitude of sinusoidal fit (2 Hz) of the membrane potential, and the spike vectorial sum during sensory stimulation, respectively. DC of membrane potential and spiking response were defined as the average membrane potential during sensory stimulation above baseline (no stimulation), and the total spike number during sensory stimulation above baseline. For spiking responses, VS neurons with firing rate > 4 spike/cycle and spike AC or DC >1 spike/cycle were included in the analysis.

### Bootstrapping

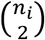 afferent pairs were counted for VS neuron *i* with *n*_*i*_ distinct afferent inputs (*n*_*i*_ ≥ 2). The same number of total afferent pairs 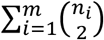 from all *m* VS neurons was randomly selected among all 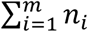 inferred afferents to determine the convergence angle or phase difference distribution by chance, and such selection was performed 5000 times to calculate mean and standard deviation.

## Acknowledgements

We thank Dr. Richard Roberts for helping set up the electrophysiology recording rig, Drs. Rebecca Callahan, Mohini Sengupta and Mr. Saul Bello Rojas for thoughtful critiques of the paper. We are grateful to Drs. Daniel Kerschensteiner and David Schoppik for insightful comments on the manuscript. We also acknowledge the Washington University Zebrafish Facility for fish care and Washington University Center for Cellular Imaging (WUCCI) for supporting the confocal imaging experiments. This work is supported by funding through the National Institute of Health (NIH) R00 DC012536 (M.W.B.), R01 DC016413 (M.W.B.), a Sloan Research Fellowship (M.W.B.), and the National BioResource Project in Japan (S.H.). M.W.B. is a Pew Biomedical Scholar and a McKnight Foundation Scholar.

## Author contributions

K.M and S.H generated the *Tg(nefma:gal4, UAS:GFP)* fish line. Z.L and M.B conceived the project. Z.L performed the electrophysiology, confocal imaging experiments and analyzed the data. T.H helped develop the deconvolution algorithm for sorting EPSC events. M.B, J.M and D.H carried out the serial section EM imaging and reconstruction. Z.L. and M.B wrote the manuscript with input from all other authors.

## References

Akrouh, A. & Kerschensteiner, D. 2013. Intersecting Circuits Generate Precisely Patterned Retinal Waves. Neuron, 79, 322–334.

Alonso, J. M. & Martinez, L. M. 1998. Functional connectivity between simple cells and complex cells in cat striate cortex. Nature Neuroscience, 1, 395–403.

Angelaki, D. E. 1992. SPATIOTEMPORAL CONVERGENCE (STC) IN OTOLITH NEURONS. Biological Cybernetics, 67, 83–96.

Angelaki, D. E., Bush, G. A. & Perachio, A. A. 1993. 2-DIMENSIONAL SPATIOTEMPORAL CODING OF LINEAR ACCELERATION IN VESTIBULAR NUCLEI NEURONS. Journal of Neuroscience, 13, 1403–1417.

Angelaki, D. E. & Cullen, K. E. 2008. Vestibular system: The many facets of a multimodal sense. Annual Review of Neuroscience, 31, 125–150.

Angelaki, D. E. & Dickman, J. D. 2000. Spatiotemporal processing of linear acceleration: Primary afferent and central vestibular neuron responses. Journal of Neurophysiology, 84, 2113–2132.

Arenz, A., Silver, R. A., Schaefer, A. T. & Margrie, T. W. 2008. The contribution of single synapses to sensory representation in vivo. Science, 321, 977–80.

Bagnall, M. W., Mcelvain, L. E., Faulstich, M. & Du Lac, S. 2008. Frequency-independent synaptic transmission supports a linear vestibular behavior. Neuron, 60, 343–52.

Bagnall, M. W. & Mclean, D. L. 2014. Modular organization of axial microcircuits in zebrafish. Science (New York, N.Y.), 343, 197–200.

Beck, A. & Teboulle, M. 2009. A Fast Iterative Shrinkage-Thresholding Algorithm for Linear Inverse Problems. Siam Journal on Imaging Sciences, 2, 183–202.

Boyle, R., Goldberg, J. M. & Highstein, S. M. 1992. Inputs from regularly and irregularly discharging vestibular nerve afferents to secondary neurons in squirrel monkey vestibular nuclei. III. Correlation with vestibulospinal and vestibuloocular output pathways. J Neurophysiol, 68, 471–84.

Boyle, R. & Johanson, C. 2003. Morphological properties of vestibulospinal neurons in primates. Annals of the New York Academy of Sciences, 1004, 183–195.

Bruno, R. M., Khatri, V., Land, P. W. & Simons, D. J. 2003. Thalamocortical angular tuning domains within individual barrels of rat somatosensory cortex. Journal of Neuroscience, 23, 9565–9574.

Budelli, R. & Macadar, O. 1979. STATO-ACOUSTIC PROPERTIES OF UTRICULAR AFFERENTS. Journal of Neurophysiology, 42, 1479–1493.

Cardona, A., Saalfeld, S., Schindelin, J., Arganda-Carreras, I., Preibisch, S., Longair, M., Tomancak, P., Hartenstein, V. & Douglas, R. J. 2012. TrakEM2 Software for Neural Circuit Reconstruction. Plos One, 7.

Chabrol, F. P., Arenz, A., Wiechert, M. T., Margrie, T. W. & Digregorio, D. A. 2015. Synaptic diversity enables temporal coding of coincident multisensory inputs in single neurons. Nat Neurosci, 18, 718–27.

Chung, J. E., Magland, J. F., Barnett, A. H., Tolosa, V. M., Tooker, A. C., Lee, K. Y., Shah, K. G., Felix, S. H., Frank, L. M. & Greengard, L. F. 2017. A Fully Automated Approach to Spike Sorting. Neuron, 95, 1381–1394 e6.

Cullen, K. E. 2019. Vestibular processing during natural self-motion: implications for perception and action. Nature Reviews Neuroscience, 20, 346–363.

Eatock, R. A. & Songer, J. E. 2011. Vestibular hair cells and afferents: two channels for head motion signals. Annu Rev Neurosci, 34, 501–34.

Ehrlich, D. E. & Schoppik, D. 2017. Control of Movement Initiation Underlies the Development of Balance. Current Biology, 27, 334–344.

Ehrlich, D. E. & Schoppik, D. 2019. A primal role for the vestibular sense in the development of coordinated locomotion. Elife, 8.

Favre-Bulle, I. A., Vanwalleghem, G., Taylor, M. A., Rubinsztein-Dunlop, H. & Scott, E. K. 2018. Cellular-Resolution Imaging of Vestibular Processing across the Larval Zebrafish Brain. Curr Biol, 28, 3711–3722 e3.

Fay, R. R. 1984. THE GOLDFISH EAR CODES THE AXIS OF ACOUSTIC PARTICLE MOTION IN 3 DIMENSIONS. Science, 225, 951–954.

Felleman, D. J. & Van Essen, D. C. 1991. Distributed Hierarchical Processing in the Primate Cerebral Cortex. Cerebral Cortex, 1, 1–47.

Fernandez, C. & Goldberg, J. M. 1976a. PHYSIOLOGY OF PERIPHERAL NEURONS INNERVATING OTOLITH ORGANS OF SQUIRREL-MONKEY.1. RESPONSE TO STATIC TILTS AND TO LONG-DURATION CENTRIFUGAL FORCE. Journal of Neurophysiology, 39, 970–984.

Fernandez, C. & Goldberg, J. M. 1976b. PHYSIOLOGY OF PERIPHERAL NEURONS INNERVATING OTOLITH ORGANS OF SQUIRREL-MONKEY.3. RESPONSE DYNAMICS. Journal of Neurophysiology, 39, 996–1008.

Fernandez, C., Goldberg, J. M. & Abend, W. K. 1972. RESPONSE TO STATIC TILTS OF PERIPHERAL NEURONS INNERVATING OTOLITH ORGANS OF SQUIRREL-MONKEY. Journal of Neurophysiology, 35, 978-+.

Goldberg, J. M., Desmadryl, G., Baird, R. A. & Fernandez, C. 1990. THE VESTIBULAR NERVE OF THE CHINCHILLA.4. DISCHARGE PROPERTIES OF UTRICULAR AFFERENTS. Journal of Neurophysiology, 63, 781–790.

Haddon, C., Mowbray, C., Whitfield, T., Jones, D., Gschmeissner, S. & Lewis, J. 1999. Hair cells without supporting cells: further studies in the ear of the zebrafish mind bomb mutant. J Neurocytol, 28, 837–50.

Hayworth, K. J., Morgan, J. L., Schalek, R., Berger, D. R., Hildebrand, D. G. C. & Lichtman, J. W. 2014. Imaging ATUM ultrathin section libraries with WaferMapper: a multi-scale approach to EM reconstruction of neural circuits. Frontiers in Neural Circuits, 8.

Hildebrand, D. G. C., Cicconet, M., Torres, R. M., Choi, W., Quan, T. M., Moon, J., Wetzel, A. W., Scott Champion, A., Graham, B. J., Randlett, O., Plummer, G. S., Portugues, R., Bianco, I. H., Saalfeld, S., Baden, A. D., Lillaney, K., Burns, R., Vogelstein, J. T., Schier, A. F., Lee, W. A., Jeong, W. K., Lichtman, J. W. & Engert, F. 2017. Whole-brain serial-section electron microscopy in larval zebrafish. Nature, 545, 345–349.

Hubel, D. H. & Wiesel, T. N. 1962. RECEPTIVE FIELDS, BINOCULAR INTERACTION AND FUNCTIONAL ARCHITECTURE IN CATS VISUAL CORTEX. Journal of Physiology-London, 160, 106-&.

Jia, H., Rochefort, N. L., Chen, X. & Konnerth, A. 2010. Dendritic organization of sensory input to cortical neurons in vivo. Nature, 464, 1307–1312.

Kimmel, C. B., Powell, S. L. & Metcalfe, W. K. 1982. Brain neurons which project to the spinal cord in young larvae of the zebrafish. The Journal of comparative neurology, 205, 112–27.

Kimura, Y., Hisano, Y., Kawahara, A. & Higashijima, S. 2014. Efficient generation of knock-in transgenic zebrafish carrying reporter/driver genes by CRISPR/Cas9-mediated genome engineering. Scientific Reports, 4.

Kodama, T., Gittis, A., Shin, M., Kelleher, K., Kolkman, K., Mcelvain, L., Lam, M. & Du Lac, S. 2020. Graded co-expression of ion channel, neurofilament, and synaptic genes in fast-spiking vestibular nucleus neurons. J Neurosci.

Korn, H., Sotelo, C. & Bennett, M. V. L. 1977. The lateral vestibular nucleus of the toadfish Opsanus tau: Ultrastructural and electrophysiological observations with special reference to electrotonic transmission. Neuroscience.

Laurens, J., Liu, S., Yu, X. J., Chan, R., Dickman, D., Deangelis, G. C. & Angelaki, D. E. 2017. Transformation of spatiotemporal dynamics in the macaque vestibular system from otolith afferents to cortex. Elife, 6.

Lavzin, M., Rapoport, S., Polsky, A., Garion, L. & Schiller, J. 2012. Nonlinear dendritic processing determines angular tuning of barrel cortex neurons in vivo. Nature, 490, 397–401.

Lecun, Y., Bengio, Y. & Hinton, G. 2015. Deep learning. Nature, 521, 436–444.

Lu, Z., Xu, Z. & Buchser, W. J. 2004. Coding of acoustic particle motion by utricular fibers in the sleeper goby, Dormitator latifrons. Journal of Comparative Physiology a-Neuroethology Sensory Neural and Behavioral Physiology, 190, 923–938.

Maruska, K. P. & Mensinger, A. F. 2015. Directional sound sensitivity in utricular afferents in the toadfish Opsanus tau. Journal of Experimental Biology, 218, 1759–1766.

Mcelvain, L. E., Faulstich, M., Jeanne, J. M., Moore, J. D. & Du Lac, S. 2015. Implementation of linear sensory signaling via multiple coordinated mechanisms at central vestibular nerve synapses. Neuron, 85, 1132–44.

Mclean, D. L. & Fetcho, J. R. 2009. Spinal Interneurons Differentiate Sequentially from Those Driving the Fastest Swimming Movements in Larval Zebrafish to Those Driving the Slowest Ones. Journal of Neuroscience, 29, 13566–13577.

Migault, G., Van Der Plas, T. L., Trentesaux, H., Panier, T., Candelier, R., Proville, R., Englitz, B., Debregeas, G. & Bormuth, V. 2018. Whole-Brain Calcium Imaging during Physiological Vestibular Stimulation in Larval Zebrafish. Curr Biol, 28, 3723–3735 e6.

Mo, W., Chen, F. Y., Nechiporuk, A. & Nicolson, T. 2010. Quantification of vestibular-induced eye movements in zebrafish larvae. Bmc Neuroscience, 11.

Nagy, J. I., Bautista, W., Blakley, B. & Rash, J. E. 2013. Morphologically mixed chemical-electrical synapses formed by primary afferents in rodent vestibular nuclei as revealed by immunofluorescence detection of connexin36 and vesicular glutamate transporter-1. Neuroscience, 252, 468–88.

Neuhuber, W. L. & Zenker, W. 1989. CENTRAL DISTRIBUTION OF CERVICAL PRIMARY AFFERENTS IN THE RAT, WITH EMPHASIS ON PROPRIOCEPTIVE PROJECTIONS TO VESTIBULAR, PERIHYPOGLOSSAL, AND UPPER THORACIC SPINAL NUCLEI. Journal of Comparative Neurology, 280, 231–253.

Newlands, S. D. & Perachio, A. A. 2003. Central projections of the vestibular nerve: a review and single fiber study in the Mongolian gerbil. Brain Research Bulletin, 60, 475–495.

Orlovsky, G. N. 1972. ACTIVITY OF VESTIBULOSPINAL NEURONS DURING LOCOMOTION. Brain Research, 46, 85-&.

Petersen, C. C. H. 2007. The functional organization of the barrel cortex. Neuron, 56, 339–355.

Peterson, B. W. 1970. Distribution of neural responses to tilting within vestibular nuclei of the cat. J Neurophysiol, 33, 750–67.

Platt, C. 1977. HAIR CELL DISTRIBUTION AND ORIENTATION IN GOLDFISH OTOLITH ORGANS. Journal of Comparative Neurology, 172, 283–297.

Priebe, N. J. & Ferster, D. 2012. Mechanisms of Neuronal Computation in Mammalian Visual Cortex. Neuron, 75, 194–208.

Priebe, N. J., Mechler, F., Carandini, M. & Ferster, D. 2004. The contribution of spike threshold to the dichotomy of cortical simple and complex cells. Nature Neuroscience, 7, 1113–1122.

Riley, B. B. & Moorman, S. J. 2000. Development of utricular otoliths, but not saccular otoliths, is necessary for vestibular function and survival in zebrafish. J Neurobiol, 43, 329–37.

Roberts, R., Elsner, J. & Bagnall, M. W. 2017. Delayed Otolith Development Does Not Impair Vestibular Circuit Formation in Zebrafish. J Assoc Res Otolaryngol, 18, 415–425.

Roy, N. C., Bessaih, T. & Contreras, D. 2011. Comprehensive mapping of whisker-evoked responses reveals broad, sharply tuned thalamocortical input to layer 4 of barrel cortex. Journal of Neurophysiology, 105, 2421–2437.

Sarkisian, V. H. 2000. Input-output relations of Deiters’ lateral vestibulospinal neurons with different structures of the brain. Archives Italiennes De Biologie, 138, 295–353.

Schoppik, D., Bianco, I. H., Prober, D. A., Douglass, A. D., Robson, D. N., Li, J. M. B., Greenwood, J. S. F., Soucy, E., Engert, F. & Schier, A. F. 2017. Gaze-Stabilizing Central Vestibular Neurons Project Asymmetrically to Extraocular Motoneuron Pools. J Neurosci, 37, 11353–11365.

Schor, R. H., Miller, A. D. & Tomko, D. L. 1984. Responses to head tilt in cat central vestibular neurons. I. Direction of maximum sensitivity. J Neurophysiol, 51, 136–46.

Sherman, S. M. 2005. Thalamic relays and cortical functioning. In: Casagrande, V. A., Guillery, R. W. & Sherman, S. M. (eds.) Cortical Function: a View from the Thalamus.

Skottun, B. C., De Valois, R. L., Grosof, D. H., Movshon, J. A., Albrecht, D. G. & Bonds, A. B. 1991. Classifying simple and complex cells on the basis of response modulation. Vision Res, 31, 1079–86.

Smear, M. C., Tao, H. Z. W., Staub, W., Orger, M. B., Gosse, N. J., Liu, Y., Takahashi, K., Poo, M. M. & Baier, H. 2007. Vesicular glutamate transport at a central synapse limits the acuity of visual perception in zebrafish. Neuron, 53, 65–77.

Straka, H. & Dieringer, N. 1996. Uncrossed disynaptic inhibition of second-order vestibular neurons and its interaction with monosynaptic excitation from vestibular nerve afferent fibers in the frog. Journal of Neurophysiology, 76, 3087–3101.

Tomko, D. L., Peterka, R. J. & Schor, R. H. 1981. RESPONSES TO HEAD TILT IN CAT 8TH NERVE AFFERENTS. Experimental Brain Research, 41, 216–221.

Turecek, J., Jackman, S. L. & Regehr, W. G. 2017. Synaptotagmin 7 confers frequency invariance onto specialized depressing synapses. Nature, 551, 503–506.

Usrey, W. M., Reppas, J. B. & Reid, R. C. 1999. Specificity and strength of retinogeniculate connections. Journal of Neurophysiology, 82, 3527–3540.

Voesenek, C. J., Pieters, R. P. M. & Van Leeuwen, J. L. 2016. Automated Reconstruction of Three-Dimensional Fish Motion, Forces, and Torques. Plos One, 11.

Wilent, W. B. & Contreras, D. 2005. Dynamics of excitation and inhibition underlying stimulus selectivity in rat somatosensory cortex. Nature Neuroscience, 8, 1364–1370.

Witts, E. C. & Murray, A. J. 2019. Vestibulospinal contributions to mammalian locomotion. Current Opinion in Physiology, 8.

