## Supplemental Figures and Methods for "Central vestibular tuning arises from patterned convergence of otolith afferents"

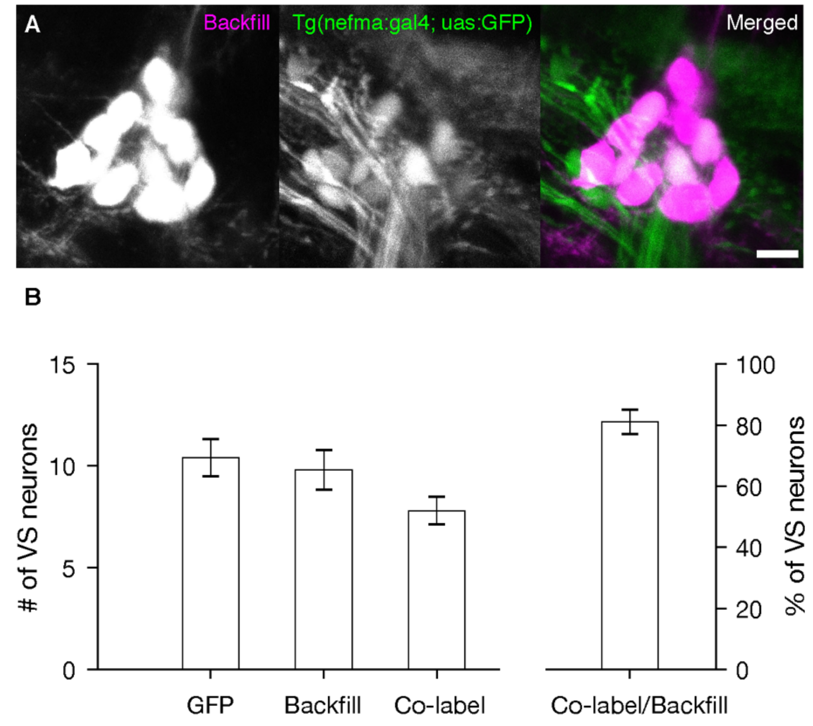


Figure S1
A. An example of labeled VS neurons by dye backfilling (magenta) from spinal cord in the Tg(nefma:gal4; uas:GFP) fishline. Scale bar: 5 $\mu$m
B. Number of labeled VS neurons (mean $\pm$ sem) by GFP, by dye backfill from the spinal cord, and co-labeled with both on one side of the brain in the larval zebrafish (5-6 dpf). Right, percentage of VS neurons identified by backfill that are also expressing GFP

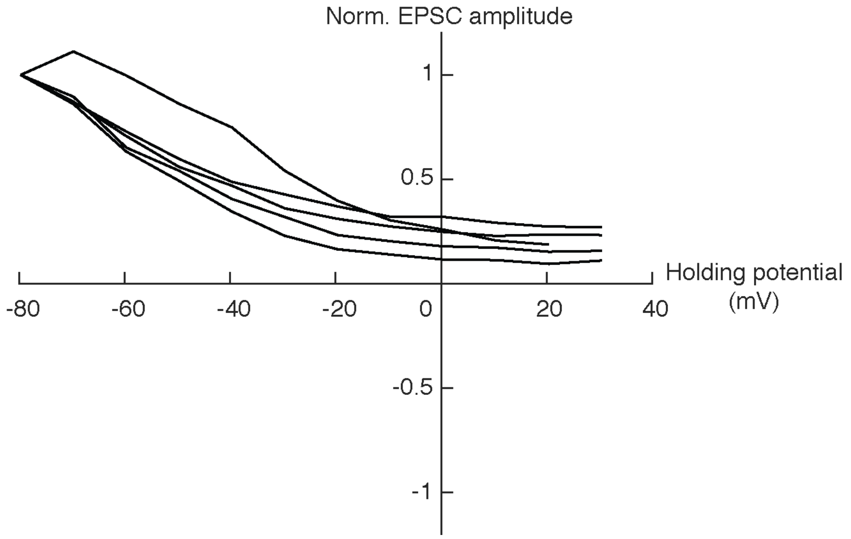


Figure S2
Normalized I-V relationship for the early EPSC component evoked by electrical stimulation on otolith afferents, normalized to EPSC amplitude at -80 mV, n=5. Note that early EPSCs do not reverse at 0 mV, consistent with electrical identity. The late EPSC (chemical) component was typically too small to measure at higher holding potential, but in some instances could be seen to reverse, consistent with chemical identity [data not shown].

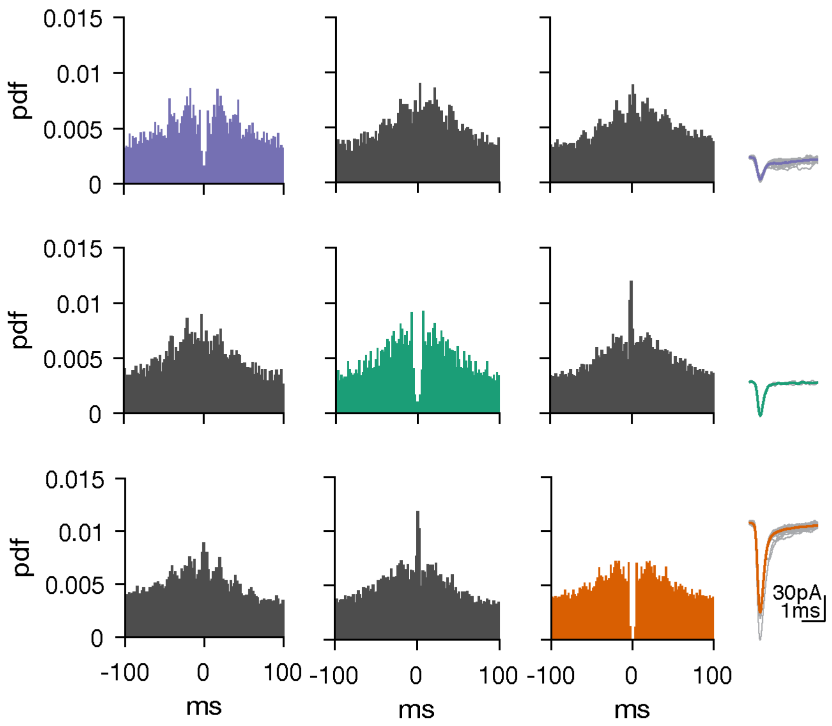


Figure S3
Auto- and cross-correlograms of three distinct EPSC clusters in one VS neuron. Note that refractory periods only appear in auto-correlograms, not in cross-correlograms.

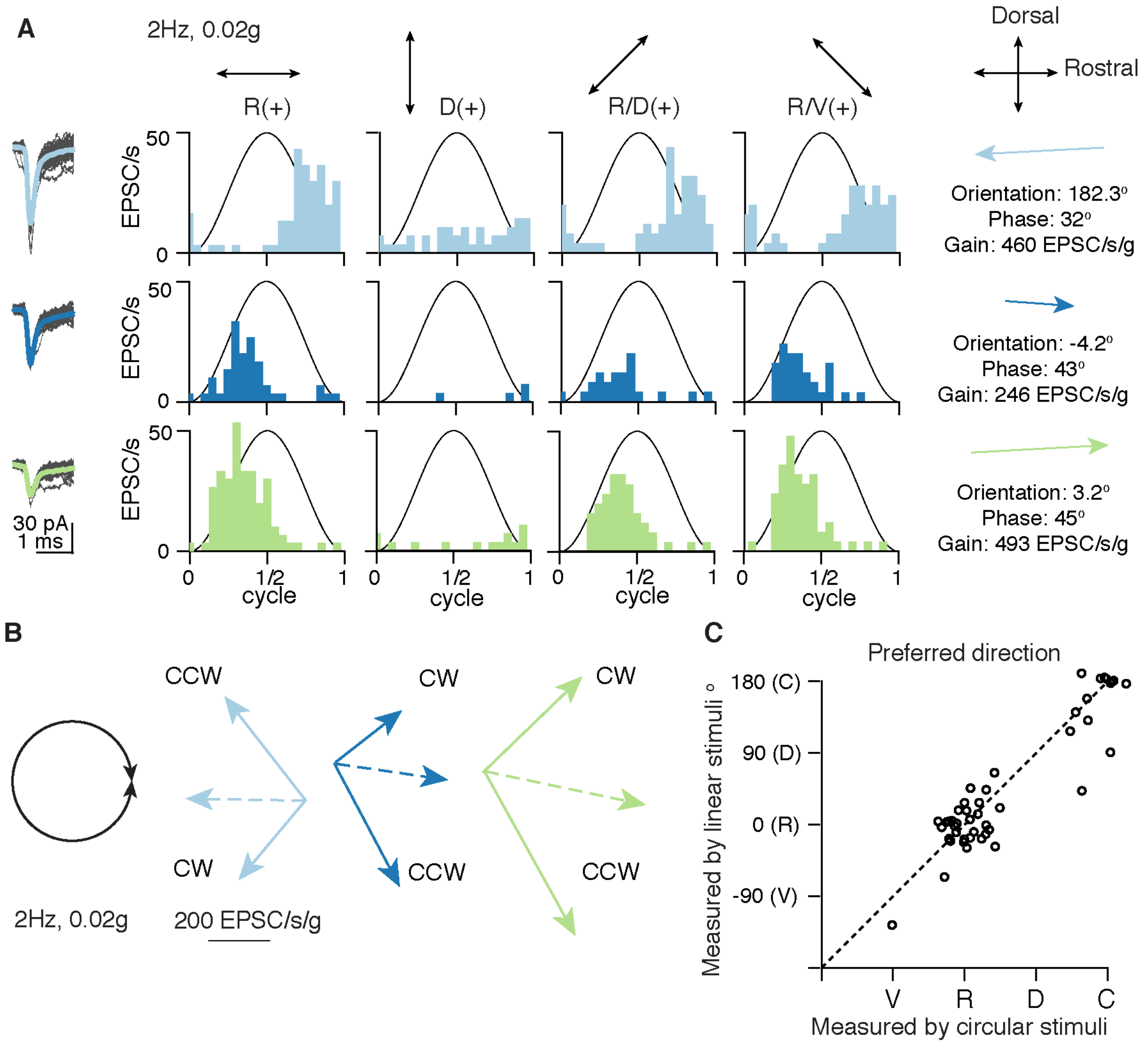


Figure S4

A. Response vectors of the three afferents converging onto one VS neuron, quantified in the same way as Fig. 5B

B. Response vectors of the same three afferents, measured with clockwise (CW) and counterclockwise (CCW) circular stimuli (left). Solid vectors represent the measured response with CW and CCW stimuli. The dashed vectors is the vector sum, an estimate of preferred tuning direction and gain as measured by circular stimuli

C. Preferred tuning direction measured with 4-axis linear stimuli was high correlated to preferred tuning direction measured with circular stimuli across the population of inferred otolith afferents (n=46), dashed: line of unity.

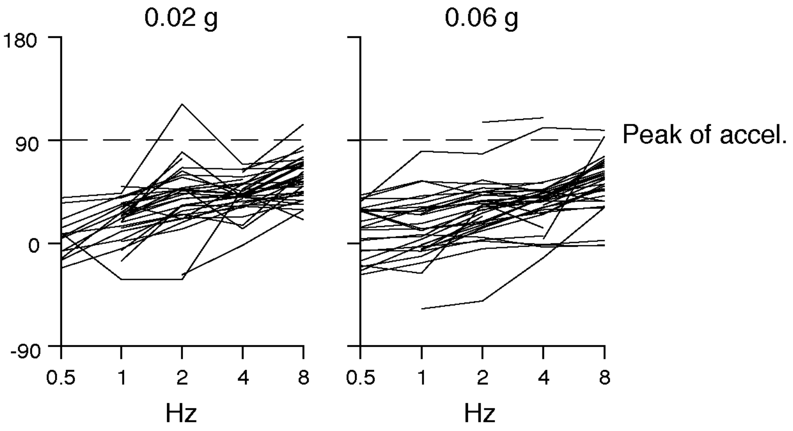


Figure S5
Phase lead relative to peak of acceleration for all otolith afferents combined. Dashed line indicates peak of acceleration towards preferred direction (either rostral or caudal).

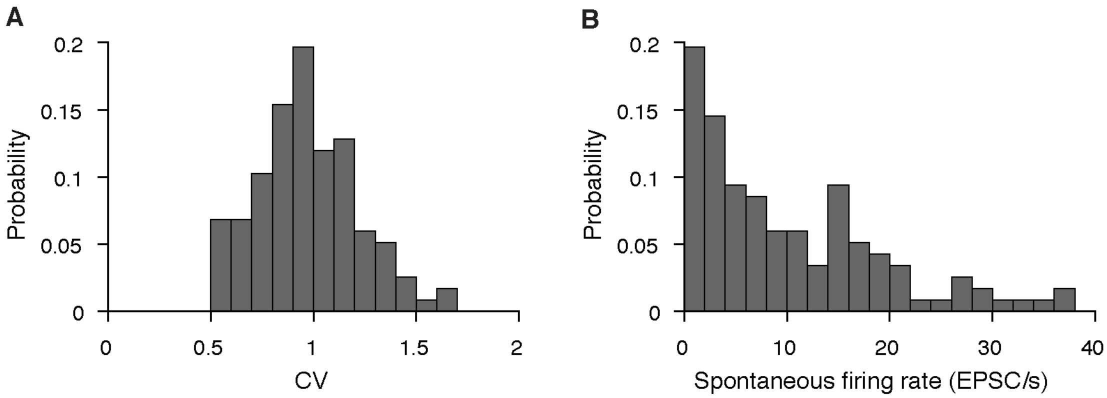


Figure S6
Distribution of CV and spontaneous firing rate of inferred otolith afferents in the larval zebrafish. (117 afferents)

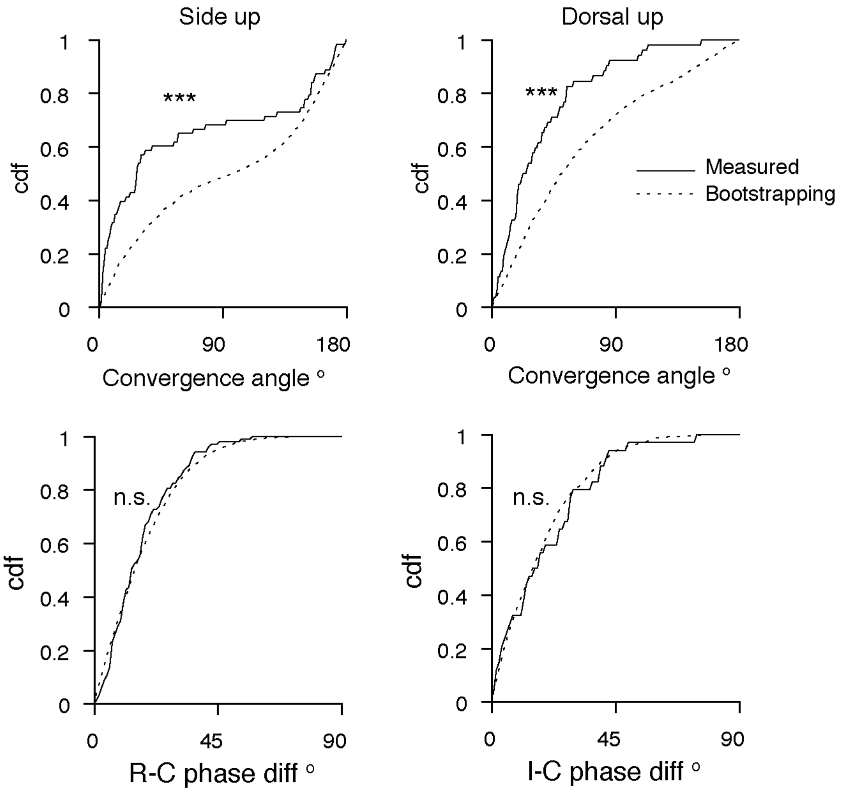


Figure. S7

Cumulative distribution function of convergence angle and phase difference, in measured (solid) and randomly generated (dashed) afferent pairs. Kolmogorov-Smirnov test, p = 1.3e-5 (convergence angle, side up), 3.5e-5 (convergence angle, dorsal up), 0.46 (R-C phase difference), 0.65 (I-C phase difference)

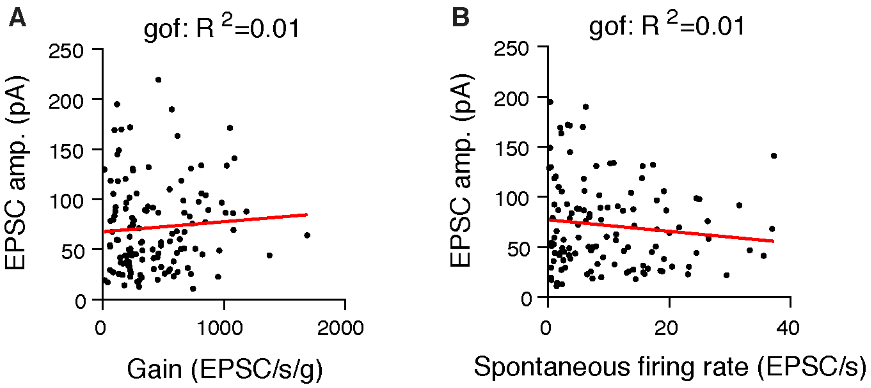


Figure S8
A. No correlation between tuning gain and EPSC amplitudes of inferred afferents (n=129), red: linear fit, gof: goodness of fit.
B. No correlation between spontaneous firing rate and EPSC amplitudes of inferred afferents (n=120), red: linear fit, gof: goodness of fit.

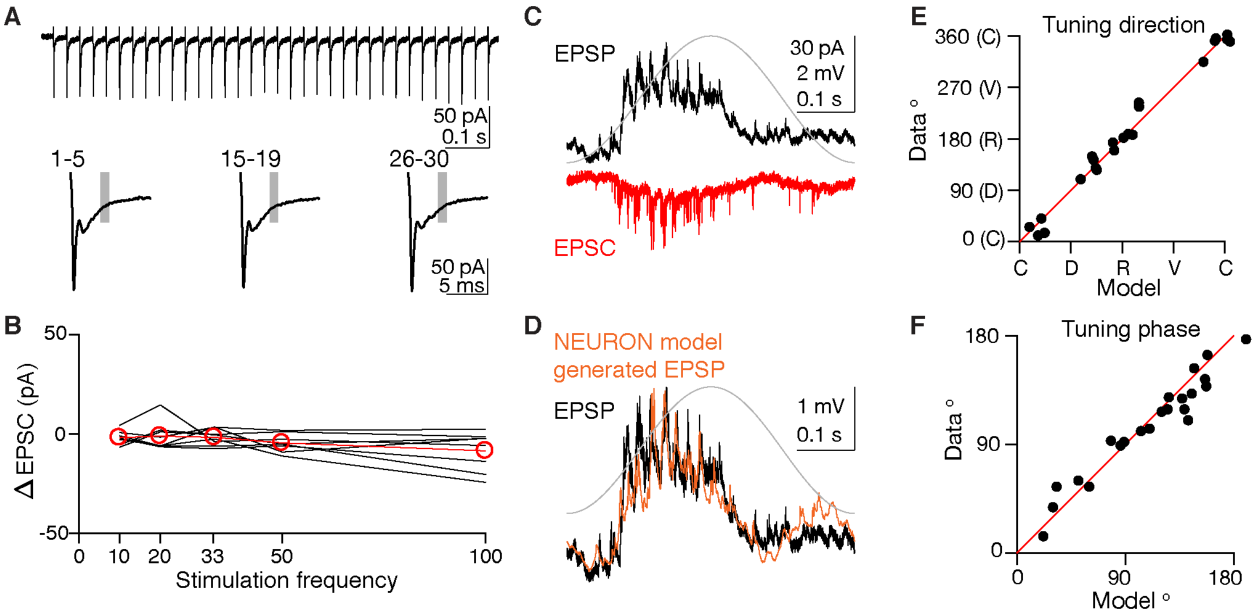


Figure S9: Excitatory afferent inputs are sufficient to explain tuning of VS neurons

A. No polysynaptic EPSCs is evoked by electrically stimulating afferents. Top, EPSCs evoked by a train (33 Hz) of electrical pulses on otolith afferents. Bottom, average traces of EPSCs at the beginning (1-5), middle (15-19) and end (26-30) during the train of electric pulses. Shaded area, 4-5 ms (estimated latency of polysynaptic EPSC) after the onset of the pulse. No additional EPSCs observed in the shaded area for all three average traces.

B. No polysynaptic EPSCs is evoked regardless of stimulation frequency on otolith afferents, n=8. ΔEPSC, difference of the average response in the shaded area (4-5 ms) to the last 5 pulses versus the first 5 pulses.

C. Comparison of EPSC (red) and EPSP (black) responses recorded sequentially in the same VS neuron, to 2 Hz, 0.02 g translational movement on the rostral(+)-caudal(-) axis.

D. Comparison of measured EPSP response (black) and model generated EPSP (Orange)

E. Tuning direction of model generated outputs is consistent with recorded EPSPs. Each dot represents the maximum tuning direction of one VS neuron, n=22.

F. Tuning phase of model generated outputs is consistent with recorded EPSPs. Each dot represents the phase of one VS neuron in the maximum direction, n=22.

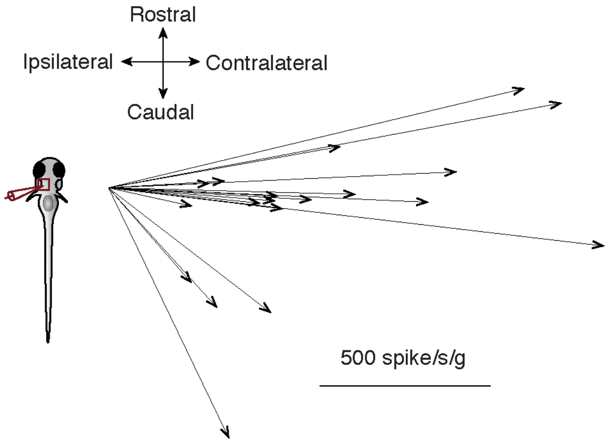


Figure. S10
Maximum direction of spiking tuning from VS neurons (n=19) in Fig. 9. Fish were oriented dorsal up.

Supplementary methods

I-V curve measurement: CsMeO4 internal solution ([in mM] CsMeSO3 122, QX314-Cl 1, TEA-Cl 1, MgCl2 3, HEPES 10, EGTA 10, Na2-ATP 4) was used to measure the reversal potential of evoked EPSCs. In the voltage clamp mode, evoked EPSCs at different holding membrane potential (-80-40 mV) were recorded. The amplitude and sign change of EPSCs were determined to plot the I-V curve. Liquid junctional potential was calculated to adjust the measured potential.

Circular stimulation: clockwise (CW) and anticlockwise (CCW) 2 Hz circular movements with direction-varying acceleration of 0.02 g, were applied at the end of the recording to measure the dynamic neural response. CW and CCW stimuli have 90° phase difference in acceleration between X and Y axes.

EPSP Modeling: Computational modeling was carried out in NEURON 7.6 (Hines and Carnevale, 1997). Because the goal was to test whether excitatory synaptic inputs were sufficient to explain the observed subthreshold tuning, only passive conductances were implemented. For each recorded neuron, input resistance measured by small hyperpolarizing steps was scaled by a fixed factor to define gLeak_hh, which was the only parameter adjusted in the model. EPSCs recorded during sensory stimulation were fed back into the model via a GClamp (dynamic clamp) mechanism (Bagnall et al., 2011) with a reversal potential of +40 mV. The resulting modeled EPSPs were then analyzed for direction and phase dynamics and compared to the same analysis on actual recorded EPSPs from the same neuron. All model code is available on https://github.com/bagnall-lab/VS_project

Reference:

BAGNALL, M. W., HULL, C., BUSHONG, E. A., ELLISMAN, M. H. & SCANZIANI, M. 2011. Multiple Clusters of Release Sites Formed by Individual Thalamic Afferents onto Cortical Interneurons Ensure Reliable Transmission. *Neuron,* 71**,** 180-194.

HINES, M. L. & CARNEVALE, N. T. 1997. The NEURON simulation environment. *Neural Computation,* 9**,** 1179-1209.
